# Characterization of the Mechanisms Underlying Sulfasalazine-Induced Ferroptotic Cell Death: Role of Protein Disulfide Isomerase-Mediated NOS Activation and NO Accumulation

**DOI:** 10.1101/2025.04.03.647136

**Authors:** Yi-Chen Jia, Jia-Ling Zhong, Xiangyu Hao, Bao Ting Zhu

## Abstract

Sulfasalazine (SAS), a clinically-utilized anti-inflammatory drug, has been shown to induce ferroptosis by inhibiting system Xc^−^ activity, thereby causing cellular glutathione depletion. Recently, it was shown that protein disulfide isomerase (PDI) is an upstream mediator of oxidative cell death (oxytosis/ferroptosis) induced by glutamate, erastin, RSL3 and SAS. The present study aims to further characterize the detailed biochemical and cellular mechanisms of SAS-induced ferroptosis in two cell lines, *i.e.*, H9C2 rat cardiomyocytes and BRL-3A rat hepatocytes, focusing on elucidating the critical role of PDI in mediating SAS-induced toxicity. We find that SAS can induce ferroptosis in H9C2 and BRL-3A cells, which is accompanied by a sequential increase in the buildup of cellular nitric oxide (NO), reactive oxygen species (ROS) and lipid-ROS. SAS activates PDI-mediated dimerization of the inducible NO synthase (iNOS) and cellular NO accumulation, and these effects are followed by ROS and lipid-ROS accumulation. Furthermore, SAS markedly upregulates the expression of iNOS protein in these cells. Knockdown of PDI or pharmacological inhibition of its catalytic activity each effectively suppresses SAS-induced iNOS dimerization, along with abrogation of SAS-induced accumulation of NO, ROS and lipid-ROS, and prevention of ferroptosis. On the other hand, PDI activation through the use of TrxR1 inhibitors sensitizes these cells to SAS-induced ferroptosis. These experimental findings provide further experimental support for a pivotal role of PDI in SAS-induced cytotoxicity through the activation of the PDI–NOS–NO axis, which then leads to cellular ROS and lipid-ROS accumulation, and ultimately induction of oxidative cell death.

## INTRODUCTION

Sulfasalazine (SAS), a well-established drug for treating inflammatory disorders such as Crohn’s disease and ulcerative colitis (1–3), has recently been identified as a potent inducer of ferroptosis (4). SAS inhibits the cystine/glutamate antiporter system Xc^−^ (5, 6), which is essential for cystine uptake and intracellular glutathione (GSH) synthesis (7). By depleting GSH, SAS disrupts the cellular redox balance and triggers oxidative stress, culminating in ferroptotic cell death. Studies have shown that SAS can induce ferroptosis in a number of cell lines (8–15), and SAS can enhance the efficacy of chemotherapeutic agents by increasing oxidative damage (5).

Ferroptosis is a regulated cell death often associated with accumulation of reactive oxygen species (ROS) (16). Unlike apoptosis or necrosis, ferroptosis has notable morphological and biochemical features, such as reduced mitochondrial size, increased membrane density, and GSH deficiency-associated oxidative damage to cellular lipids (17). Central to this process is the depletion of cellular GSH and inactivation of glutathione peroxidase 4 (GPX4), and these changes disrupt a cell’s ability to neutralize reactive lipid peroxides (lipid-ROS), subsequently leading to cell death (18).

Protein disulfide isomerase (PDI or PDIA1) is a well-known member of the thioredoxin superfamily, and is primarily localized in the endoplasmic reticulum (19). Functionally, PDI serves as a dithiol/disulfide oxidoreductase, and facilitates protein folding by catalyzing the isomerization of the intra- and intermolecular disulfide bonds (20). Recent studies from our laboratory have shown that PDI is involved in mediating chemically-induced, glutathione (GSH) depletion-associated ferroptosis (21–23). Specifically, GSH depletion leads to PDI oxidation and activates its catalytic activity for nitric oxide synthase (NOS) dimerization via disulfide bond formation, which subsequently results in the buildup of cellular nitric oxide (NO) and ROS/lipid-ROS and mitochondrial ROS, and ultimately ferroptotic cell death (21–23). Notably, inhibition of PDI function has been shown to block NOS dimerization and NO accumulation and thus effectively rescues cells from chemically-induced ferroptosis (21–23). These findings position PDI as a pivotal upstream mediator of ferroptosis, linking GSH depletion to oxidative cell death pathways.

Recently, we have shown that PDI is involved in mediating SAS-induced ferroptosis using the immortalized HT22 mouse hippocampal neurons as an *in-vitro* model (15). In the present study, we seek to further characterize the detailed cellular and biochemical mechanisms of SAS-induced ferroptotic cell death using two rat cell lines, namely, the H9C2 cardiomyocytes and BRL-3A hepatocytes. These two cell lines are selected as *in-vitro* models for study because earlier studies have shown that they are sensitive to erastin- and RSL3-induced ferroptosis in culture (22–27). In Moreover, recently we have shown that PDI is involved in mediating erastin-induced ferroptotic cell death in BRL-3A rat hepatocytes (25). Confirming our recent observations (15), the results of our present study further demonstrate that PDI plays a pivotal role in mediating SAS-induced ferroptosis in both H9C2 cardiomyocytes and BRL-3A hepatocytes, which involves PDI-mediated NOS dimerization, accumulation of NO, ROS and lipid-ROS, and ultimately ferroptotic cell death.

## MATERIALS AND METHODS

### Chemicals and reagents

Sulfasalazine (SAS, #HY-14655), BAZ and ferrostatin-1 (Fer-1, #HY-100579) were obtained from MedChemExpress (Monmouth Junction, NJ, USA), *N*-acetyl-*L*-cysteine (NAC, #A8199), cystamine dihydrochloride (cystamine, #C121509), thiazolyl blue tetrazolium bromide (MTT, #T818538) from MACKLIN (Shanghai, China); carboxy-PTIO (cPTIO, #S1547), diaminofluorescein-FM diacetate (DAF-FM-DA, #S0019) and 2’,7’-dichlorodihydrofluorescein diacetate (DCFH-DA, #S0033S) from Beyotime Biotechnology (Shanghai, China); BODIPY-581/591-C11 (#D3861) from ThermoFisher Scientific (Waltham, Massachusetts, USA); EN460 (#M07515) from BioLab (Beijing, China).

The anti-iNOS antibody (#ab15323) was obtained from Abcam (Cambridge, UK), the anti-PDI antibody (#3501S) and anti-β-actin (#4970) antibodies were purchased from Cell Signaling Technology (Boston, Massachusetts, USA); goat anti-mouse IgG and goat anti-rabbit IgG conjugated to horseradish peroxidase (#7074S) were from Santa Cruz Biotechnology (Santa Cruz, CA, USA) and used as secondary antibodies. Most of the other chemicals were obtained from Sigma-Aldrich (Saint Louis, Missouri, USA).

### Cell culture and viability assay

The rat H9C2 cardiomyocytes and BRL-3A hepatocytes were obtained from the Cell Bank of the Chinese Academy of Sciences (Shanghai, China), and were maintained in DMEM medium supplemented with 10% (*v*/*v*) FBS (fetal bovine serum; ThermoFisher, Waltham, MA, USA) and antibiotics (100 U/mL penicillin + 100 μg/mL streptomycin; Sigma-Aldrich) at 37℃ under 5% CO_2_. The cells were passaged or used in experiments upon reaching approximately 80% confluence, and they were usually under 25 passages. Authentication was performed by STR profiling and routine mycoplasma testing.

For cell viability assays, cells were first seeded in 96-well plates (at 2,000 cells/well), and treated with different drugs as indicated. MTT (0.5 mg/mL) was added to each well and incubated for 3 h at 37°C under 5% CO_2_. DMSO was added afterwards to dissolve the MTT formazan, and absorbance was measured at 560 nm wavelength with a BioTek microplate reader (BioTek, Winooski, VT, USA).

### siRNA transfection

The procedures of siRNA transfection of cultured cells were described in our earlier study (24, 25). Briefly, 24 h after seeding, siRNAs (at 60 nM) for targeted genes (PDI, iNOS, nNOS or TrxR1) were transfected using Lipofectamine RNA iMAX (Invitrogen). Forty-eight h after siRNA transfection, cells were treated with the selected drugs, and subsequently processed for cell viability determination, immunoblotting and fluorescence imaging.

### Measurement of cellular NO and ROS levels by fluorescence microscopy

Cells were plated in 24-well plates at a density of 5 ⨯ 10^4^ per well, and then given different drug treatments as indicated. For fluorescence microscopy analysis of cellular NO and total ROS, the cells were first washed twice with HBSS and then incubated with DAF-FM-DA (at 5 μM for staining of NO) and DCFH-DA (at 5 μM for staining of ROS), respectively, in 200 μL DMEM (serum-free and phenol red-free) for 20 min at 37°C under 5% CO_2_. Following three washes with HBSS, fluorescence images were taken with an AXIO fluorescence microscope (Carl Zeiss Corporation, Germany).

### Confocal microscopy

For visualization of the subcellular distribution of cellular lipid-ROS, cells were seeded at a density of 5 ⨯ 10^4^ per well on coverslips placed inside the 24-well plates. Twenty-four h later, cells were treated with drugs as indicated. Coverslips were then washed in HBSS and incubated in HBSS containing BODIPY-581/591-C11 (5 μM) for 20 min at 37°C under 5% CO_2_. Coverslips were then mounted on microscope slides for visualization. Slides were imaged using a LSM 900 confocal laser scanning microscope (Carl Zeiss), and images were analyzed with the Zen software (Carl Zeiss).

### Flow cytometry

For flow cytometric analysis of cellular levels of NO, ROS and lipid-ROS, the cells were plated in 6-well plates at a density of 15 ×10^4^ cells/well for 24 h before treatment with different drugs. Afterwards, cells were trypsinized, collected and suspended in phosphate-buffered saline (PBS). Cells were then centrifuged, and the resulting cell pellets were resuspended in DMEM (free of phenol red and serum) containing DAF-FM-DA (5 μM), DCFH-DA (5 μM) and BODIPY-581/591-C11 (5 μM), respectively. After a 20-min incubation at 37°C, the cells were washed three times with HBSS to remove any remaining fluorescent dyes. Cellular NO, ROS and lipid-ROS were measured using a flow cytometer (Beckman Coulter, Brea, CA, USA), and the data were analyzed using the FlowJo software (FlowJo, LLC, Ashland, USA).

### Western blot analysis

Following treatment of the cells with SAS for varying durations as indicated, the cells were collected by trypsinization and centrifugation, and then lysed on ice for 30 min in a RIPA buffer containing 1% protease inhibitor cocktail. Protein concentrations were determined using the BCA Assay Kit (ThermoFisher). For total iNOS protein analysis (including both monomeric and dimeric forms of iNOS), samples were heated at 95°C for 5 min with a reducing buffer before loaded onto the electrophoresis gel. To analyze the monomeric and dimeric forms of iNOS, the samples were prepared in a non-reducing buffer and were not heated, and the temperature of the gel was maintained below 15°C during electrophoresis. The proteins were separated using 10% agarose gel (for total iNOS) or 6% agarose gel (for monomeric and dimeric iNOS), and then transferred to PVDF membranes. The membranes were incubated for 1 h in 5% skim milk at room temperature, and then incubated with the primary antibody overnight. Afterwards, the membranes were washed three times with TBST at room temperature (10 min each time). The membranes were then incubated with the secondary antibody for 1 h at room temperature and washed with TBST three times before visualization. Desired protein bands were visualized using the chemiluminescence imaging system (Tanon 5200, Shanghai, China). The experiments were repeated multiple times to confirm the observations. Representative blots from one representative experiment are shown.

### Statistical analysis

In this study, the quantitative experiments were repeated multiple times to confirm the experimental observations. Statistical analyses were carried out using one-way ANOVA followed by Dunnett’s post-hoc tests for multiple comparisons (GraphPad Prism 7.0 software; GraphPad Software, La Jolla, CA). The data were presented as the mean ±standard deviation (S.D.), typically obtained from a representative experiment with multiple replicates. *P* < 0.05 (* or ^#^) and *P* < 0.01 (** or ^##^) indicate statistically significant and very significant differences, respectively. In most cases, * and ** denotes comparisons for statistical significance between the control group (cells treated with vehicle only) and the cells treated with SAS, whereas ^#^ and ^##^ denote comparisons between the cells treated with SAS and the cells jointly treated with SAS + another compound. For Western blot quantification, one representative data set is shown.

## RESULTS

### Induction of ferroptotic cell death by SAS in H9C2 and BRL-3A cells

The cytotoxicity of SAS in H9C2 rat cardiomyocytes was evaluated by exposing the cells to increasing concentrations (0.125, 0.25, 0.5, 1 and 2 mM) of SAS over a 24-h period, with cell viability assessed using the MTT assay. SAS elicited cytotoxicity in H9C2 cells in a time- and concentration-dependent manner (**Fig. 1A, 1B**), with an *IC*_50_ of ∼0.28 mM. SAS-induced cell death was further verified with Calcein-AM/PI double staining (**Fig. 1C**). Morphological changes of cells resembling ferroptosis were noted at 6‒8 h after SAS exposure (**Supplementary Fig. S1A**). At the concentration of 0.5 mM SAS, cell survival rate dropped to ∼20%, making this concentration ideal for further studying the protective effects of selected compounds against SAS-induced cytotoxicity.

**Figure 1.**
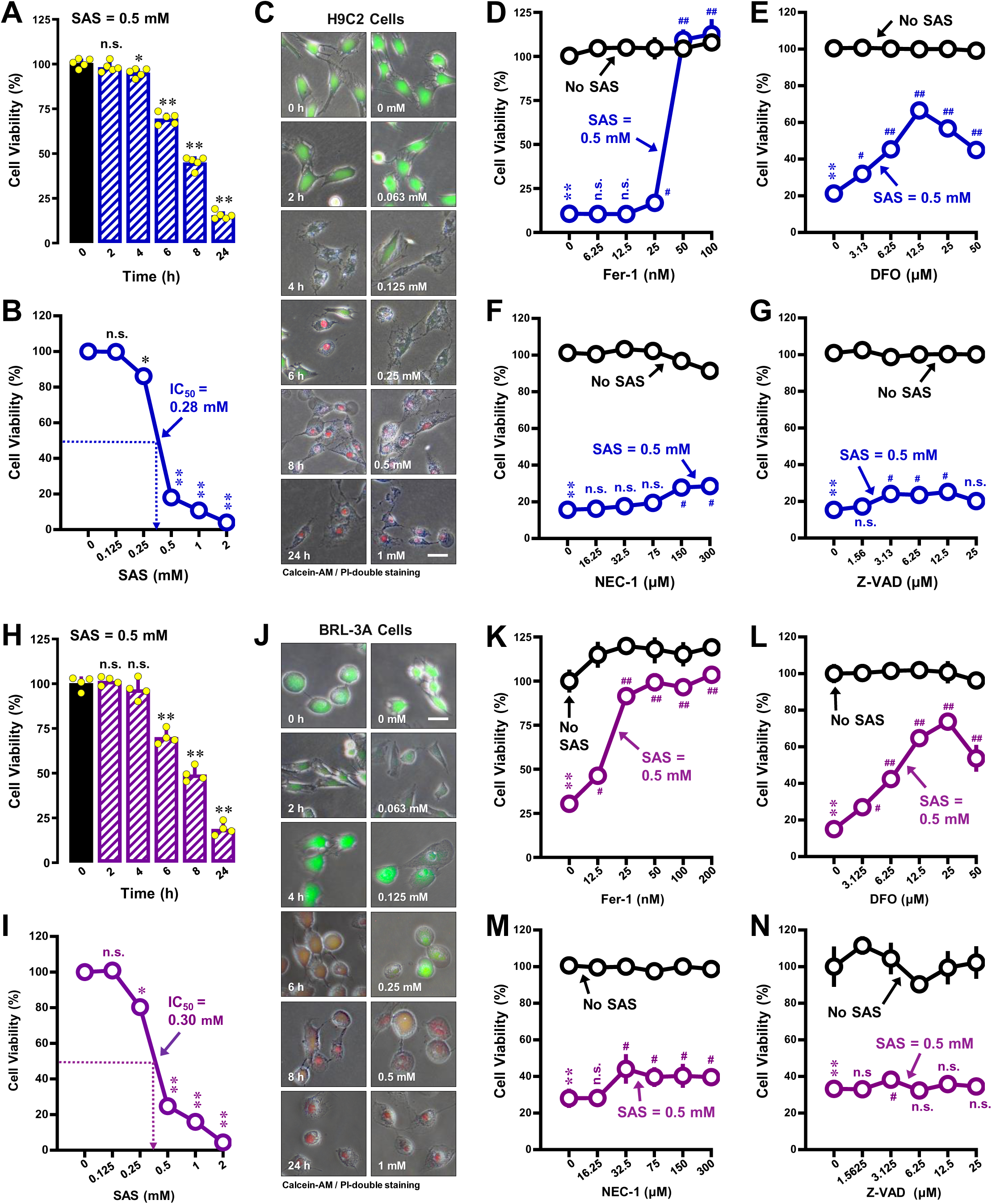
Induction of ferroptotic cell death by SAS in H9C2 and BRL-3A cells. **A, B, C, H, I, J.** Time- and does-dependent induction of cell death by SAS in H9C2 (**A, B, C**) and BRL-3A (**H, I, J**) cells. Cells were treated with different concentrations of SAS for 24 h or treated with 0.4 mM SAS for 0, 2, 4, 6, 8 and 24 h. Cell viability was determined by MTT assay (**A, B, H, I**; n = 5) or by fluorescence microscopy (**C, J**) following Calcein-AM/PI staining (20×, scale bar = 100 μm). **D, E, F, G.** Effect of Fer-1 (**D**), DFO (**E**), NEC-1 (**F**) and z-VAD-FMK (**G**) on SAS-induced death in H9C2 cells. Cells were treated with SAS (0.5 mM) ±Fer-1, DFO, z-VAD-FMK or NEC-1 for 24 h, and then subject to MTT assay (n = 5). **K, L, M, N.** Effect of Fer-1 (**K**), DFO (**L**), NEC-1 (**M**) and z-VAD-FMK (**N**) on SAS-induced death in BRL-3A cells. Cells were treated with SAS (0.5 mM) ±Fer-1, DFO, z-VAD-FMK or NEC-1 for 24 h, and then subjected to MTT assay (n = 5). Quantitative data are presented as mean ±SD. * or ^#^ *P* < 0.05; ** or ^##^ *P* < 0.01; n.s., not significant.

To ascertain whether SAS-induced cell death in H9C2 cells qualifies as ferroptosis, we investigated the protective effects of Fer-1, deferoxamine (DFO) and necrostatin-1 (Nec-1) on SAS-induced cytotoxicity. Fer-1, a known antioxidant for lipid-ROS and a prototypical ferroptosis inhibitor (28), completely rescued SAS-induced cell death when present at 50‒ 100 μM (**Fig. 1D**). Likewise, DFO, an iron chelator known for its ferroptosis-rescuing activity (29, 30), also provided substantial protection (**Fig. 1E**). In comparison, necrostatin-1 (NEC-1), an inhibitor of necrosis (31, 32), and z-VAD-FMK, a pan-caspase inhibitor (33, 34), did not exhibit a similar protection against SAS-induced cell death (**Fig. 1F, 1G**).

SAS-induced ferroptotic cell death was also evaluated in BRL-3A rat hepatocytes. SAS exhibited a time- and concentration-dependent cytotoxicity in these cells (**Fig. 1H, 1I** for MTT assay; **Fig. 1J** for Calcein-AM/PI staining), with an *IC*_50_ of ∼0.30 mM. Morphological changes consistent with ferroptosis were observed around 8 h post SAS exposure (**Supplementary Fig. S1B**). Consistent with the induction of ferroptosis by SAS in BRL-3A cells, Fer-1 and DFO mitigated SAS-induced cell death (**Fig. 1K, 1L**), but NEC-1 and z-VAD-FMK did not exert a similar protection against SAS-induced cell death (**Fig. 1M, 1N**). Collectively, these findings strongly indicate that SAS triggers a form of cell death that closely aligns with ferroptosis in both H9C2 and BRL-3A cells.

### Effects of SAS on cellular NO, ROS and lipid-ROS accumulation

Our recent research has revealed that the accumulation of NO, ROS and lipid-ROS plays a pivotal role in the induction of ferroptotic cell death by erastin (22, 23) and RSL3 (24). To explore whether similar accumulations also contribute to SAS-induced ferroptosis, we first investigated the time- and concentration-dependent changes in cellular NO, ROS and lipid-ROS levels in H9C2 cells. A small increase in cellular NO levels (using DAF-FM-DA as a probe) was observed between 2 and 4 h following SAS exposure, with a more pronounced increase occurring between 6 and 8 h (**Fig. 2A, 2C**). As expected, SAS also elicited a concentration-dependent increase in cellular NO levels (**Fig. 2D**).

**Figure 2.**
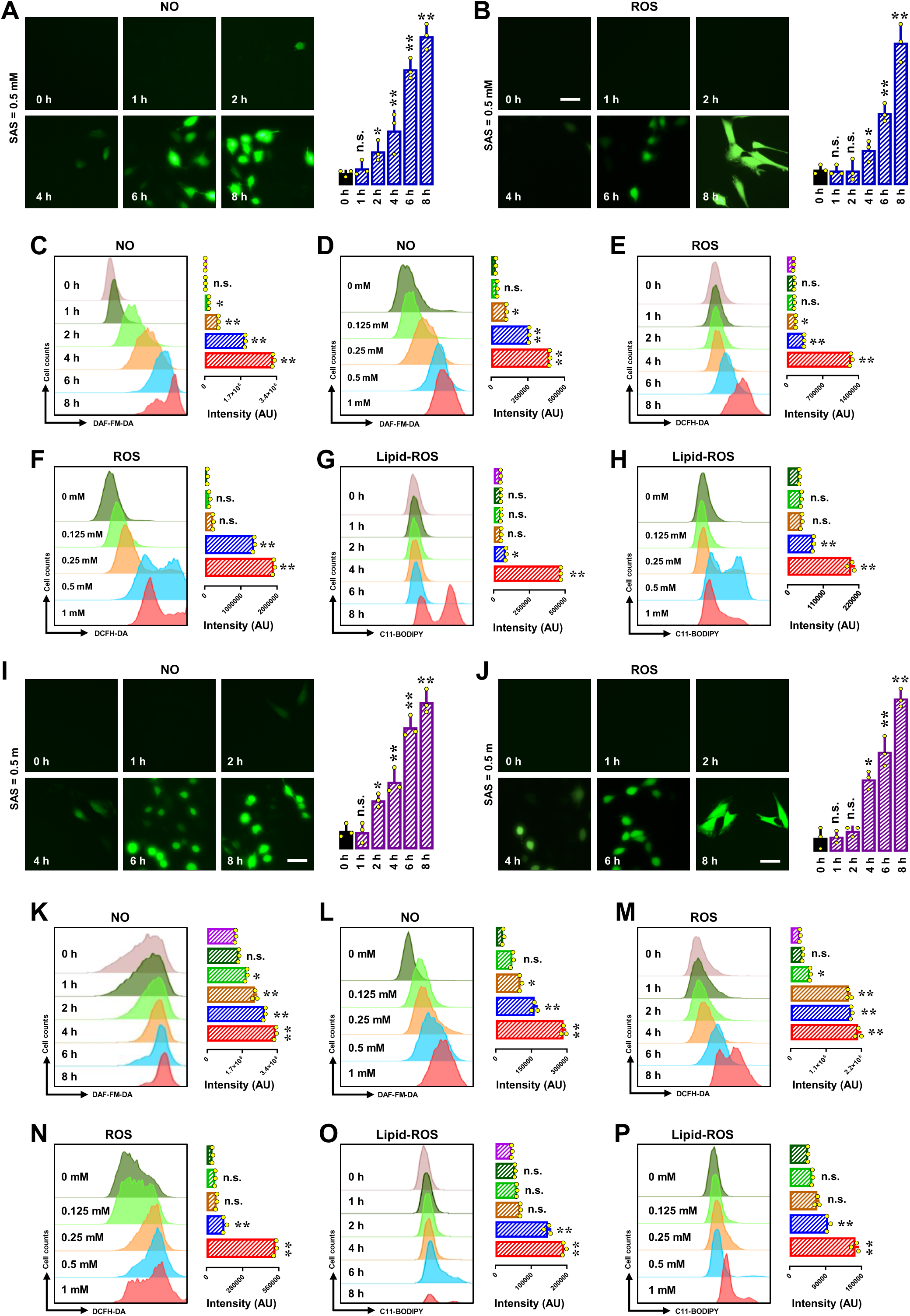
Time- and concentration-dependent change in cellular NO, ROS and lipid-ROS levels in SAS-treated H9C2 and BRL-3A cells. **A‒H.** Time and concentration dependency of SAS-induced accumulation of NO (**A, C, D**), ROS (**B, E, F**) and lipid-ROS (**G, H**) in H9C2 cells. Cells were treated with 0.5 mM SAS for varying durations as specified (for time dependency) or treated with different concentrations of SAS as indicated for 8 h (for concentration dependency). Then the cells were subjected to fluorescence microscopy (scale bar = 100 μm) or analytical flow cytometry. For the fluorescence microscopy data (**A, B**), the quantitative intensity values are shown in the right panels; similarly, the intensity values for flow cytometry data (**C–H**) are also shown in the right panels (n = 3). **I‒P.** Time and concentration dependency of SAS-induced accumulation of NO (**I, K, L**), ROS (**J, M, N**) and lipid-ROS (**O, P**) in BRL-3A cells. The treatment conditions, measurements and figure annotations are the same as described above for panels **A‒H**. Quantitative data are presented as mean ±S.D. \**P* < 0.05; ** *P* < 0.01 *vs* the control group; n.s., not significant.

Similarly, SAS exposure caused a time-dependent increase in cellular ROS levels (using DCFH-DA as a probe; **Fig. 2B, 2E**). An initial small increase in cellular ROS was detectable at 4–6 h post-exposure, peaking at 8 h (**Fig. 2B, 2E**). Additionally, there was a concentration-dependent increase in ROS levels in SAS-treated cells (**Fig. 2F**). The rise in cellular ROS levels appeared to be behind in time the buildup of cellular NO.

Lipid-ROS accumulation is a defining feature of chemically-induced ferroptosis (35, 36). In this study, we examined changes in cellular lipid-ROS levels in SAS-treated H9C2 cells (using C11-BODIPY as a probe). We observed that SAS elicited time- and concentration-dependent increases in cellular lipid-ROS levels in these cells (**Fig. 2G, 2H**).

We also investigated the time- and concentration-dependent changes in cellular NO, ROS and lipid-ROS levels in BRL-3A cells (**Fig. 2I–2P**). SAS caused a modest increase in NO levels between 2 and 4 h following SAS exposure, and a more pronounced increase occurring between 6 and 8 h (**Fig. 2I, 2K**). As expected, SAS elicited a concentration-dependent increase in cellular NO levels (**Fig. 2L**).

Similarly, SAS exposure also elicited time- and concentration-dependent increases in cellular ROS levels (**Fig. 2J, 2M** for time dependency; **Fig. 2N** for concentration dependency) and cellular lipid-ROS levels (**Fig. 2O, 2P**) in BRL-3A cells.

Collectively, these results demonstrate that SAS elicits a sequential and time-dependent increase in cellular NO, ROS and lipid-ROS levels in both H9C2 and BRL-3A cells. It is evident that SAS-induced NO accumulation precedes the cellular accumulations of ROS and lipid-ROS.

### Effect of NO, ROS and lipid-ROS scavengers on SAS-induced cell death

To determine the contribution of cellular NO, ROS and lipid-ROS accumulation in mediating SAS-induced ferroptosis in H9C2 cells, representative scavengers of NO, ROS and lipid-ROS were employed to evaluate their modulating effects on SAS-induced ferroptosis by measuring the change in cell viability and cellular NO, ROS and lipid-ROS levels.

#### cPTIO

cPTIO is an NO scavenger commonly used in cell culture studies (37). To explore whether NO accumulation leads to ROS/lipid-ROS buildup and subsequent cell death, we examined the impact of cPTIO on SAS-induced ferroptosis in H9C2 cells. Our findings revealed that cPTIO partially mitigated SAS-induced cell death (**Fig. 3A** for MTT assay; **Fig. 3B** for Calcein-AM/PI double staining). The modest cytoprotective effect of cPTIO may be related to its own cytotoxicity as this chemical can produce cytotoxic NO_2_ during its scavenging of NO (38). We also investigated the effect of cPTIO on the accumulation of cellular NO, ROS and lipid-ROS. SAS-induced NO accumulation was partially reduced by co-treatment with 100 μM cPTIO (**Fig. 3C, 3D**); in comparison, the SAS-induced buildup of cellular ROS (**Fig. 3E, 3F**) and lipid-ROS (**Fig. 3G**) was more strongly reduced by cPTIO. Joint treatment of cells with cPTIO also abrogated the SAS-induced increases in cellular NOS protein levels, including total, dimer and monomer forms (**Fig. 3O, left panel**).

**Figure 3.**
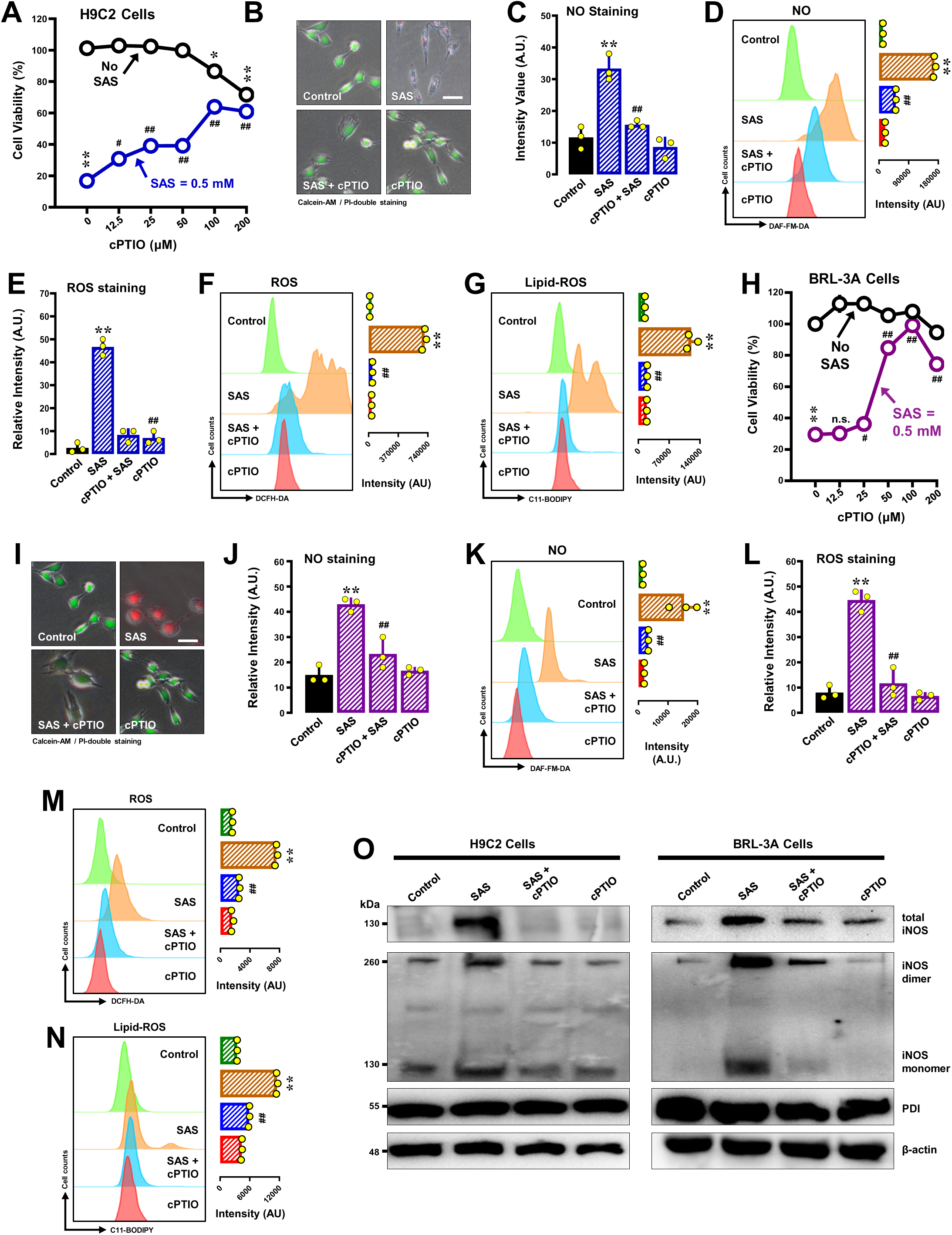
Effects of cPTIO on SAS-induced cytotoxicity, accumulation of NO, ROS and lipid-ROS, and iNOS dimerization in H9C2 and BRL-3A cells. **A, B, H, I.** Protective effect of cPTIO on SAS-induced cytotoxicity in H9C2 (**A, B**) and BRL-3A (**H, I**) cells. In **A** and **H**, cells were treated with SAS (0.4 mM) ±cPTIO (12.5, 25, 50, 100 and 200 µM) for 24 h, and then subjected to MTT assay (n = 4). In **B** and **I**, cells were treated with SAS (0.4 mM) ±cPTIO (50 μM) for 24 h, and then subjected to fluorescence microscopy following Calcein-AM/PI staining (green for live cells, and red for dead cells; scale bar = 100 μm). **C, D, E, F, G.** Abrogation by cPTIO of SAS-induced accumulation of cellular NO (**C, D**), ROS (**E, F**) and lipid-ROS (**G**) in H9C2 cells. Cells were treated with SAS (0.5 mM) ± cPTIO (100 μM) for 8 h, and then subjected to fluorescence microscopy (**C, E**) and analytical flow cytometry (**D, F, G**). For the fluorescence microscopy data in **C, E**, only the quantitative intensity values are shown (n = 3). For flow cytometry data (**D, F, G**), the intensity values are shown in the right panels (n = 3). **J, K, L, M, N.** Abrogation by cPTIO of SAS-induced accumulation of cellular NO (**J, K**), ROS (**L, M**) and lipid-ROS (**N**) in BRL-3A cells. Cells were treated with SAS (0.5 mM) ± cPTIO (100 μM) for 8 h, and then subjected to fluorescence microscopy (**J, L**) and analytical flow cytometry (**K, M, N**). For the fluorescence microscopy data in **J, L**, only the quantitative intensity values are shown (n = 3). For flow cytometry data (**K, M, N**), the corresponding intensity values are shown in the right panels (n = 3). **O.** Effect of cPTIO on SAS-induced changes in total, monomer and dimer iNOS protein levels in H9C2 and BRL-3A cells (Western blotting). Cells were treated with SAS (0.5 mM) ±cPTIO (100 µM) for 8 h, and then levels of the total, monomer and dimer iNOS proteins and PDI were determined by Western blotting. β-Actin was used as a loading control. Quantitative data are presented as mean ±SD. * or ^#^ *P* < 0.05; ** or ^##^ *P* < 0.01; n.s., not significant.

Similar analyses were performed in BRL-3A cells. We found that cPTIO partially mitigated SAS-induced cell death (**Fig. 3H** for MTT assay; **Fig. 3I** for Calcein-AM/PI staining). SAS-induced NO accumulation was partially reduced by co-treatment with 100 μM cPTIO (**Fig. 3J, 3K**). Interestingly, SAS-induced buildup of cellular ROS (**Fig. 3L, 3M**) and lipid-ROS (**Fig. 3N**) was more significantly reduced by cPTIO. cPTIO also reduced SAS-induced increases in total, dimer and monomer forms of the iNOS protein (**Fig. 3O, right panel**).

#### NAC

*N*-Acetyl-*L*-cysteine (NAC) is an antioxidant known for its robust protective effect against oxidative stress-associated cytotoxicity (39, 40). In H9C2 cells, joint treatment with 1 mM NAC strongly prevented SAS-induced cell death, achieving nearly complete cytoprotection when present at 1 mM (**Fig. 4A, 4B**). NAC also abrogated SAS-induced accumulation of NO (**Fig. 4C, 4D**), ROS (**Fig. 4E**) and lipid-ROS (**Fig. 4G, 4H**) in these cells, illustrating its broad-spectrum antioxidant capacity.

**Figure 4.**
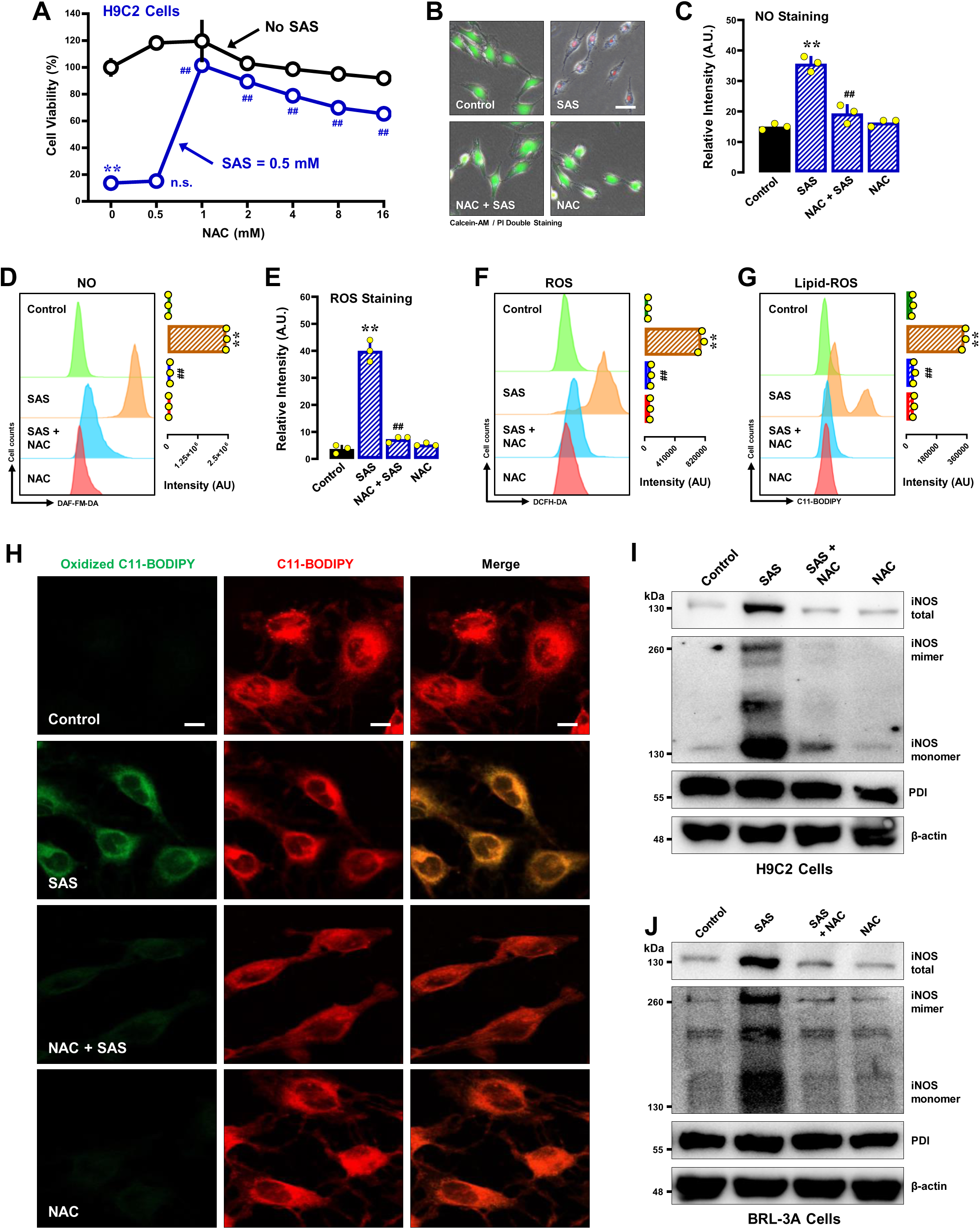
Effect of NAC on SAS-induced ferroptosis and accumulation of NO, ROS and lipid-ROS in H9C2 and BRL-3A cells. **A, B.** Protective effect of NAC against SAS-induced cytotoxicity in H9C2 cells. In **A**, cells were treated with SAS (0.5 mM) ±NAC (0.5, 1, 2, 4, 8 and 16 mM) for 24 h, and then subjected to MTT assay (n = 4). In **B**, cells were treated with SAS (0.5 mM) ±NAC (1 mM) for 24 h, and then subjected to fluorescence microscopy following Calcein-AM/PI staining (green for live cells, and red for dead cells; scale bar = 100 μm). **C, D, E, F, G, H**. Abrogation by NAC of SAS-induced accumulation of cellular NO (**C, D**), ROS (**E, F**) and lipid-ROS (**G**, **H**) in H9C2 cells. Cells were treated with SAS (0.5 mM) ± NAC (1 mM) for 8 h, and then subjected to fluorescence microscopy (**C**, **E**), flow cytometry (**D, F, G**) and confocal microscopy (**H**; scale bar = 100 μm). For the fluorescence microscopy data in **C, E**, only the quantitative intensity values are shown (n = 3). For flow cytometry data (**D**, **F**, **G**), the left panels are the histograms, and the right panels are the quantitative values (n = 3). **I, J.** Effect of NAC on SAS-induced changes in total, monomer and dimer iNOS protein levels in H9C2 (**I**) and BRL-3A (**J**) cells. Cells were treated with SAS (0.5 mM) ±NAC (1 mM) for 8 h, and then levels of the total, monomer and dimer iNOS proteins and PDI were determined by Western blotting. β-Actin was used as a loading control. Quantitative data are presented as mean ±SD. ** or ^##^ *P* < 0.01; n.s., not significant.

In BRL-3A cells, NAC also prevented SAS-induced cell death (**Supplementary Fig. S2A** for MTT assay; **Supplementary Fig. S2B** for Calcein-AM/PI staining). Moreover, NAC strongly mitigated SAS-induced accumulation of cellular NO (**Supplementary Fig. S2C**), ROS (**Supplementary Fig. S2D**) and lipid-ROS (**Supplementary Fig. S2E, S2F**).

In addition to its effects on oxidative markers, joint treatment with NAC suppressed SAS-induced increases in cellular levels of total iNOS and its monomeric and dimeric forms (**Fig. 4I** for H9C2 cells; **Fig. 4J** for BRL-3A cells).

#### Fer-1

Fer-1 exhibited a robust protection against SAS-induced ferroptosis at a low concentration (50 nM), exhibiting nearly 100% protection in H9C2 cells (**Fig. 5A** for MTT assay; **Fig. 5B** for Calcein-AM/PI double staining). Fer-1 effectively abrogated SAS-induced accumulation of cellular NO, ROS and lipid-ROS (**Fig. 5C 5D** for NO; **Fig. 5E, 5F** for ROS; **Fig. 5G, 5H** for lipid-ROS).

**Figure 5.**
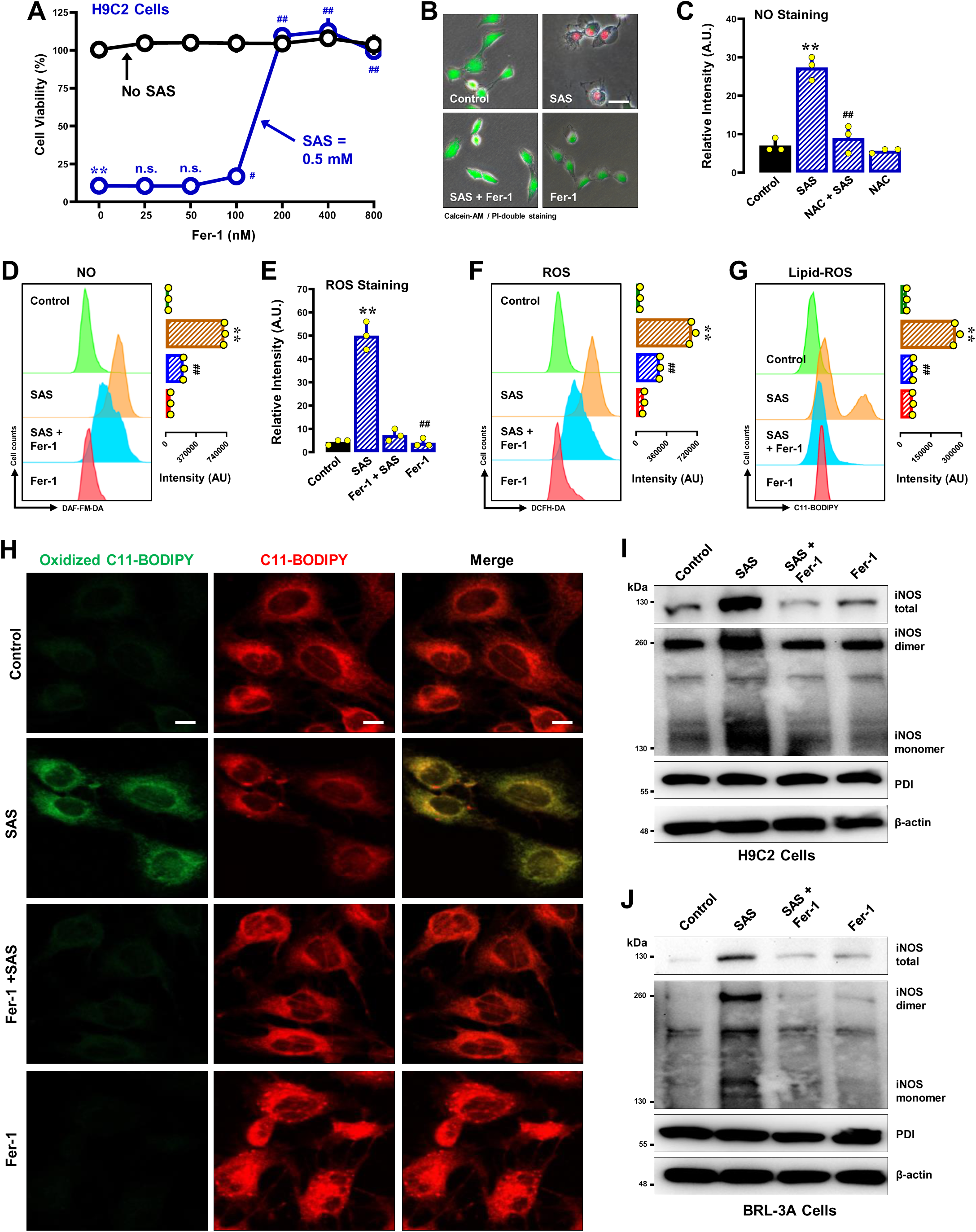
Effect of Fer-1 on SAS-induced ferroptosis and accumulation of NO, ROS and lipid-ROS in H9C2 and BRL-3A cells. **A, B.** Protective effect of NAC against SAS-induced cytotoxicity. In **A**, H9C2 cells were treated with SAS (0.5 mM) ±Fer-1 (25, 50, 100, 200, 400 and 800 nM) for 24 h, and then cell viability was determined by MTT assay (n = 4). In **B**, cells were treated with SAS (0.5 mM) ±Fer-1 (200 nM) for 24 h, and then fluorescent images of Calcein-AM/PI-stained cells were captured (green for live cells, and red for dead cells; scale bar = 100 μm). **C, D, E, F, G, H**. Abrogation by Fer-1 of SAS-induced accumulation of cellular NO (**C, D**), ROS (**E, F**) and lipid-ROS (**G, H**) in H9C2 cells. Cells were treated with SAS (0.5 mM) ± Fer-1 (200 nM) for 8 h, and then subjected to fluorescence microscopy (**C, E**), flow cytometry (**D, F, G**) and confocal microscopy (**H**; scale bar = 100 μm). For the fluorescence microscopy data in **C, E**, only the quantitative intensity values are shown (n = 3). For flow cytometry data (**D, F, G**), the left panels are the histograms, and the right panels are the quantitative values (n = 3). **I, J.** Effect of Fer-1 on SAS-induced changes in total, monomer and dimer iNOS protein levels in H9C2 (**I**) and BRL-3A (**J**) cells. Cells were treated with SAS (0.5 mM) ±Fer-1 (200 nM) for 8 h, and then levels of the total, monomer and dimer iNOS proteins and PDI were determined by Western blotting. β-Actin was used as a loading control. Quantitative data are presented as mean ±SD. * or ^#^ *P* < 0.05; ** or ^##^ *P* < 0.01; n.s., not significant.

Similarly, Fer-1 was also strongly protected BRL-3A cells against SAS-induced ferroptosis (**Supplementary Fig. S3A** for MTT assay; **Supplementary Fig. S3B** for Calcein-AM/PI staining). Fer-1 abrogated SAS-induced accumulation of cellular NO, ROS and lipid-ROS (**Supplementary Fig. S3C** for NO; **Fig. S3D** for ROS; **Fig. S3E, S3F** for lipid-ROS). Additionally, Fer-1 abrogated SAS-induced increases in total cellular iNOS protein as well as its mononer and dimer (**Fig. 5I** for H9C2 cells; **Fig. 5J** for BRL-3A cells).

**DFO.** Deferoxamine (DFO) is an iron chelator with known ferroptosis-rescuing properties (29, 30). DFO was shown to have significant protection against SAS-induced cytotoxicity, as shown by MTT assay (**Supplementary Fig. S4A** for H9C2 cells; **Supplementary Fig. S4F** for BRL-3A cells) and Calcein-AM double staining (**Supplementary Fig. S4B** for H9C2 cells; **Supplementary Fig. S4G** for BRL-3A cells). Furthermore, DFO effectively abrogated SAS-induced accumulation of cellular NO, ROS and lipid-ROS (**Supplementary Fig. S4C–S4E** for H9C2 cells; **Supplementary Fig. S4H–S4J** for BRL-3A cells).

Trolox, a compound known for its potent ROS-scavenging activity and limited NO-scavenging capability (41), provided nearly complete protection against SAS-induced cytotoxicity in H9C2 and BRL-3A cells (**Supplementary Fig. S5A** for H9C2 cells; **Supplementary Fig. S5B** for BRL-3A cells). In contrast, MitoTEMPO, a mitochondrial-targeting antioxidant and superoxide dismutase mimetic that converts mitochondrial superoxide anion to H_2_O_2_ (42), did not show any protective effect against SAS-induced cytotoxicity in these two cell lines (**Supplementary Fig. S5C** for H9C2 cells; **Supplementary Fig. S5D** for BRL-3A cells). In contrast, TEMPO exerted a partial protection against SAS-induced cytotoxicity (**Supplementary Fig. S5E** for H9C2 cells; **Supplementary Fig. S5F** for BRL-3A cells).

Together, these findings indicate that SAS induces the accumulation of cellular NO, ROS and lipid-ROS, which collectively mediate the induction of oxidative ferroptosis in both H9C2 and BRL-3A cells.

### Effect of SNP on SAS-induced ferroptotic cell death

To provide additional support for the hypothesis that in SAS-induced NO accumulation is a key event which subsequently leads to ROS/lipid-ROS accumulation, we investigated the effect of sodium nitroprusside (SNP), a known NO donor (43), on ROS and lipid-ROS accumulation as well as on ferroptotic cell death in H9C2 and BRL-3A cells that were jointly treated with SAS and SNP.

Using the MTT assay, we found that 300 μM SNP alone produced minimal cytotoxicity in H9C2 cells (**Fig. 6A**). However, co-treatment with 100 μM SNP significantly enhanced SAS-induced cytotoxicity (**Fig. 6B**). Our earlier studies have shown that SNP alone can lead to cellular accumulation of ROS and lipid-ROS (22–24). In this study, we found that joint treatment of H9C2 cells with SNP (300 μM) + SAS markedly accelerated the accumulation of cellular ROS (**Fig. 6C, 6D**) and lipid-ROS (**Fig. 6E**). Similar observations were also made in BRL-3A cells. While SNP at 300 μM produced minimal cytotoxicity in these cells (**Fig. 6F**), joint treatment with 100 μM SNP significantly enhanced SAS-induced cytotoxicity (**Fig. 6G**). In addition, joint treatment of cells with SNP (300 μM) markedly accelerated SAS-induced accumulation of cellular ROS (**Fig. 6H, 6I**) and lipid-ROS (**Fig. 6J**). These findings suggest that elevated cellular NO levels can amplify SAS-induced ferroptosis by accelerating cellular ROS/lipid-ROS accumulation.

**Figure 6.**
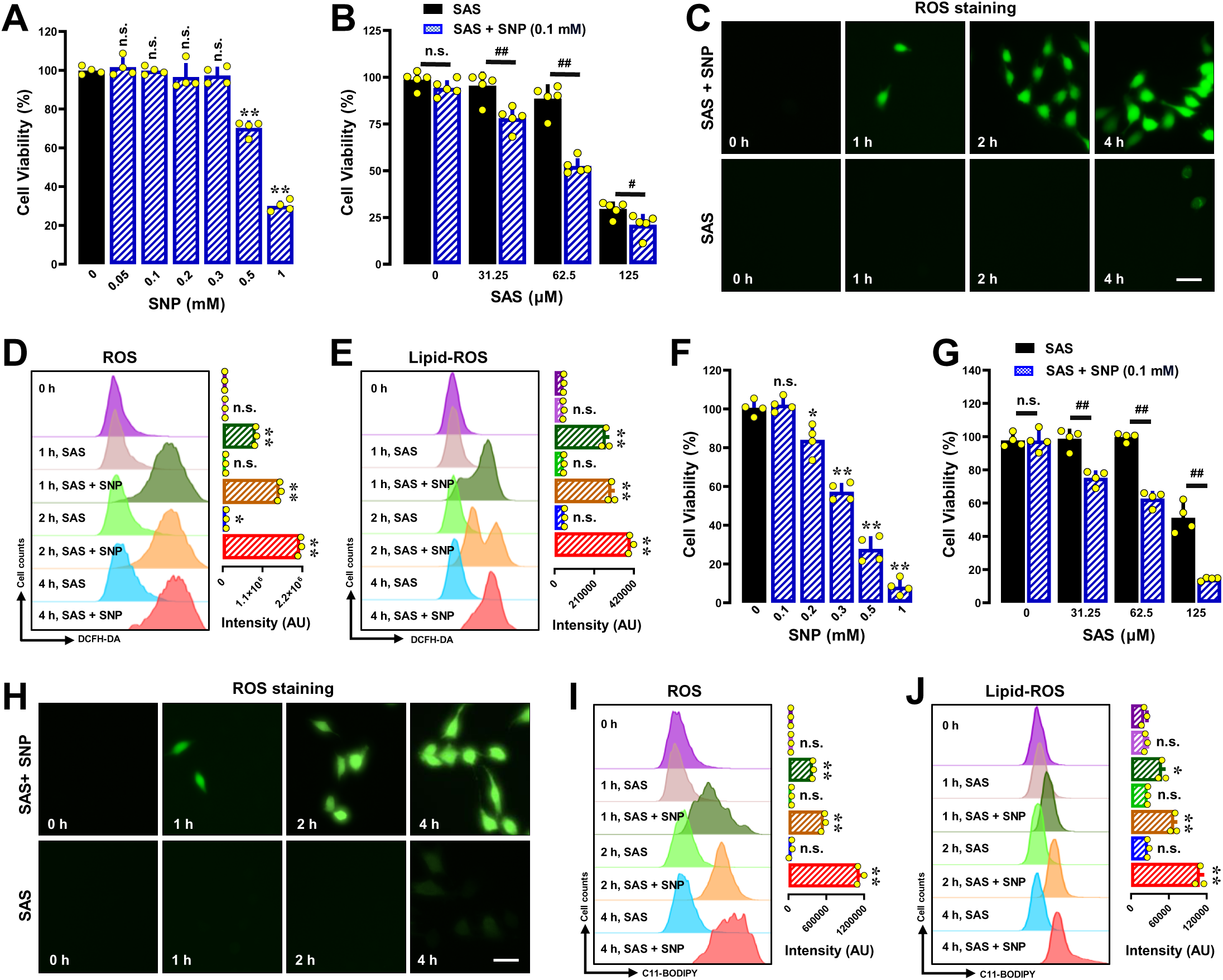
Effect of SNP on SAS-induced ferroptotic H9C2 and BRL-3A cell death. **A, C**. Concentration-dependent cytotoxicity of SNP in H9C2 (**A**) and BRL-3A (**C**) cells (MTT assay, n = 4). **F, G.** Enhancement of SAS-induced ferroptosis by SNP in H9C2 (**F**) and BRL-3A (**G**) cells. Cells were exposed to increasing concentrations of SAS ±SNP (0.1 mM) for 24 h, and cell viability was assessed using the MTT assay (n = 4). **C, D, E, H, I, J.** Time-dependent accumulation of cellular ROS (**C, D, H, I**) and lipid-ROS (**E, J**) in H9C2 and BRL-3A cells. Cells were treated with SAS (0.5 mM) ±SNP (0.1 mM) for 1, 2 and 4 h, and then subjected to fluorescence microscopy (**C, H**) or for flow cytometry analysis (**D, E, I, J**). For flow cytometry data, the left panels are histograms, and the right panels are the corresponding quantitative values (n = 3). Quantitative data are presented as mean ±SD. ** or ^##^ *P* < 0.01; n.s., not significant.

### Effect of SAS on iNOS protein levels and their dimerization

The formation of cellular NO is catalyzed by NOS using *L*-arginine as substrate (44). In this study, we evaluated the effect of SAS on the protein levels and particularly their dimers in H9C2 and BRL-3A cells. We found that iNOS and eNOS were expressed in H9C2 cells (iNOS data shown in **Fig. 7A**; eNOS data not shown), but nNOS is basically not detected (data not shown). The iNOS protein levels were increased in a concentration- and time-dependent manner, with a marked induction seen at 6 h after SAS exposure (**Fig. 7A**). In comparison, cellular levels of PDI were not significantly affected by SAS (**Fig. 7A**).

**Figure 7.**
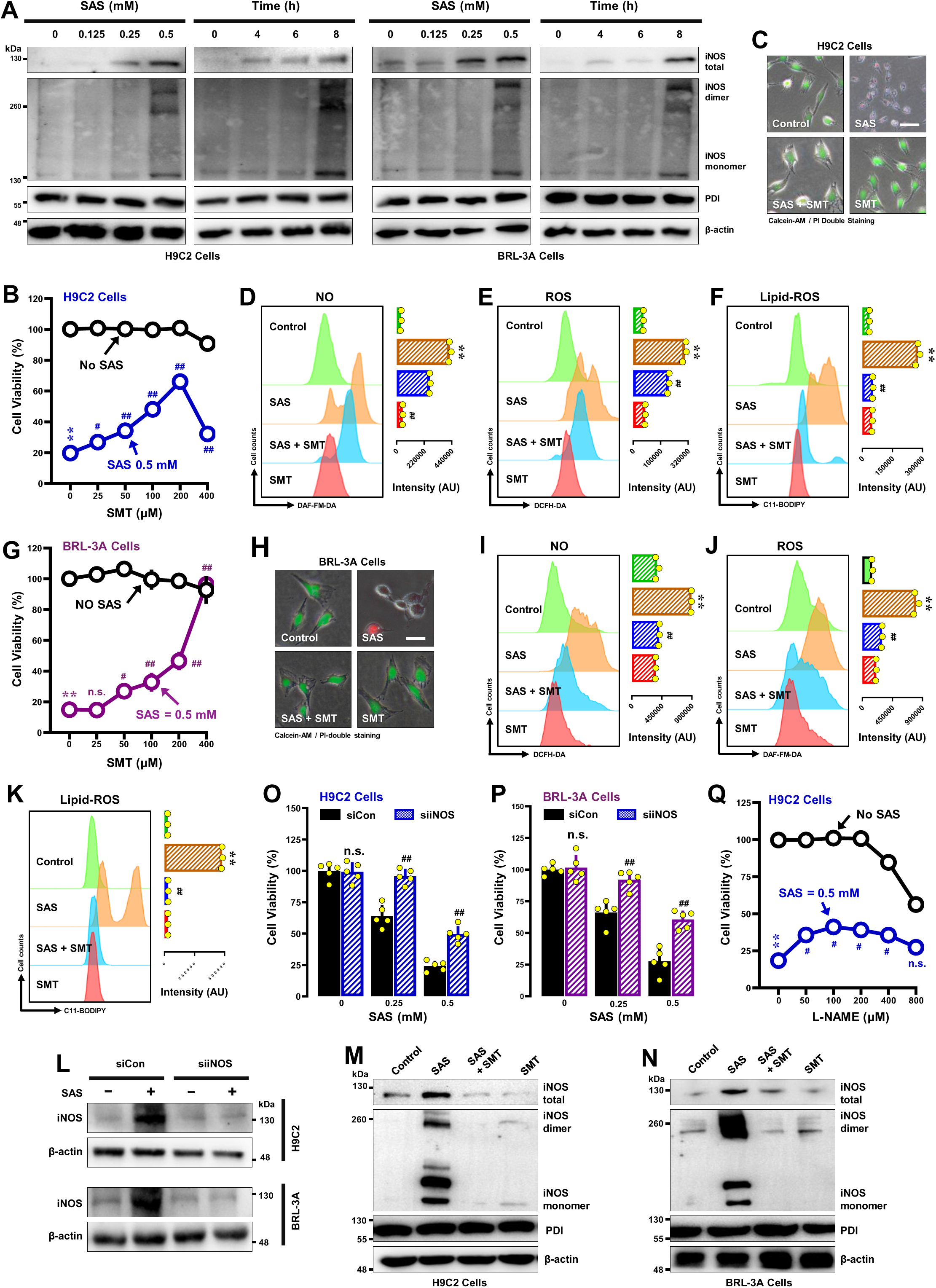
Effect of SAS on iNOS upregulation and dimer formation in H9C2 and BRL-3A cells. **A**. Concentration- and time-dependent effects of SAS on cellular levels of total, monomer and dimer iNOS proteins (Western blotting) in H9C2 (**K**) and BRL-3A (**L**) cells. Cells were treated with different concentrations of SAS for 8 h or with 0.5 mM SAS for different durations, and then cellular levels of total, monomer and dimer iNOS proteins were determined by Western blotting. Cellular PDI and β-actin levels were also determined for comparison. **B, C, G, H.** Protective effect of SMT against SAS-induced cytotoxicity in H9C2 (**B, C**) and BRL-3A cells (**G, H**). In **B** and **G**, cells were exposed to SAS (0.5 mM) ±SMT (25, 50, 100, 200 and 400 μM) for 24 h, and then subjected to MTT assay (n = 5). In **C** and **H**, cells were treated with SAS (0.5 mM) ±SMT (200 μM for H9C2 cells/400 μM for BRL-3A cells) for 24 h, and then subjected to fluorescence microscopy following Calcein-AM/PI staining (**C, H**; green color for live cells and red color for dead cells; scale bar = 100 μm). **D-F, I-K**. Abrogation by SMT of SAS-induced accumulation of cellular NO (**D, I**), ROS (**E, J**) and lipid-ROS (**F, K**) in H9C2 and BRL-3A cells. Cells were treated with SAS (0.5 mM) ± SMT (200 μM for H9C2 cells/400 μM for BRL-3A cells) for 8 h, and then subjected to flow cytometry. For flow cytometry data, the left panels are the histograms, and the right panels are the quantitative values (n = 3). **N.** Effectiveness of iNOS-siRNAs in reducing cellular iNOS protein levels in H9C2 and BRL-3A cells. Cells were transfected with iNOS-siRNAs for 48 h prior to treatment of cells with SAS (0.5 mM) for an additional 8 h. **Q.** Effect of *L*-NAME on SAS-induced cytotoxicity in H9C2 cells. Cells were treated with SAS (0.5 mM) ±*L*-NAME (25, 50, 100, 200, 400 and 800 μM) for 24 h, and then subjected to MTT assay (n = 5). **O, P.** Effect of iNOS knockdown on SAS-induced cytotoxicity in H9C2 (**O**) and BRL-3A (**P**) cells. Cells were transfected with iNOS-siRNAs 48 h prior to treatment of cells with SAS (0.5 mM) for an additional 24 h, and cell viability was determined by MTT assay (n = 5). **M, N.** Effect of SMT on SAS-induced changes in total, monomer and dimer iNOS protein levels (Western blotting). H9C2 (**M**) and BRL-3A (**N**) cells were treated with SAS (0.5 mM) ±SMT (200 μM for H9C2 cells/400 μM for BRL-3A cells) for 8 h, and then the cellular total, monomer and dimer iNOS proteins were determined by Western blotting. Quantitative data are presented as mean ±SD. * or ^#^ *P* < 0.05; ** or ^##^ *P* < 0.01; n.s., not significant.

In BRL-3A cells, only iNOS was detected, and its protein levels were similarly upregulated in a concentration- and time-dependent manner (**Fig. 7A**). nNOS and eNOS proteins were largely undetectable in these cells (data not shown). The cellular levels of the PDI protein were not significantly affected by SAS treatment in these cells (**Fig. 7A**).

#### NOS inhibitors

*S*-Methylisothiourea sulfate (SMT) is an iNOS inhibitor (45). We found that joint treatment of H9C2 cells with SAS + SMT for 24 h exerted a partial protection against SAS-induced ferroptosis (**Fig. 7B** for MTT assay; **Fig. 7C** for Calcein-AM/PI double staining). SMT partially reduced SAS-induced accumulation of cellular NO (**Fig. 7D**) and ROS (**Fig. 7E**), but strongly abrogated SAS-induced lipid-ROS accumulation (**Fig. 7F**). Similar to the effects seen in H9C2 cells, joint treatment of BRL-3A cells with SMT strongly rescued SAS-induced ferroptosis (**Fig. 7G** for MTT assay; **Fig. 7H** for Calcein-AM/PI double staining). SMT reduced SAS-induced accumulation of NO (**Fig. 7I**) and ROS (**Fig. 7J**), and more strongly abrogated SAS-induced lipid-ROS accumulation (**Fig. 7K**). In addition, SMT abrogated SAS-induced increase in total iNOS proteins and their dimer levels (**Fig. 7M** for H9C2 cells; **Fig. 7N** for BRL-3A cells). For comparison, we also examined the effect of *L*-NAME, an inhibitor of eNOS (46, 47), on SAS-induced ferroptotic cell death. We found that *L*-NAME only elicited a very small protection against SAS-induced ferroptosis (**Fig. 7Q**). Collectively, these results collectively indicate that NO accumulation resulting from the induction and activation (dimerization) of iNOS in SAS-treated cells is involved in mediating SAS-induced ferroptosis in H9C2 and BRL-3A cells.

#### iNOS knockdown

Based on the above studies with NOS inhibitors, next we employed the siRNA approach to selectively knock down iNOS expression to further evaluate its role in mediating SAS-induced ferroptosis in H9C2 and BRL-3A cells. The effectiveness of iNOS knockdown was assessed by Western blotting of total iNOS protein levels (**Fig. 7L**). Notably, knockdown of iNOS resulted in a significant protection against SAS-induced cytotoxicity in both H9C2 (**Fig. 7O**) and BRL-3A cells (**Fig. 7P**). Notably, while eNOS was also expressed in H9C2 cells, we did not use the siRNA approach to examine its role in SAS-induced ferroposis as *L*-NACK, an eNOS inhibitor (46, 47), only elicits a very small protection (see **Fig. 7Q**).

### PDI is an upstream mediator of SAS-induced ferroptosis

#### PDI knockdown

Our earlier studies have shown that the activated PDI (*i.e.*, the disulfide bonds in PDI’s catalytic sites are oxidized) mediates chemically-induced oxidative ferroptosis through its catalysis of NOS dimerization (23, 48). To determine whether the activated PDI also mediates NOS dimerization in SAS-treated cells, first we determined whether SAS can directly activate the catalytic activity of PDI in the *in-vitro* enzymatic assays. As shown in **Supplementary Fig. S6A, 6B**, the presence of SAS at 250 and 500 μM had little or no effect on the oxidative catalytic activity of PDI. In addition, the presence of SAS at these two concentrations did not affect the reductive activity of PDI (**Supplementary Fig. S6C, S6D**).

Next, we selectively knocked down PDI expression in H9C2 cells using PDI-siRNAs for 24 and 48 h. Western blotting confirmed the effectiveness of PDI knockdown at the protein level (**Fig. 8A**). Next, we assessed the cytotoxicity of SAS-treated H9C2 cells at two different concentrations (0.25 and 0.5 mM) of SAS. DPI knockdown caused significant reductions in the sensitivity of H9C2 cells to SAS-induced cytotoxicity (**Fig. 8B** for 24 h knockdown; **Fig. 8C** for 48 h knockdown). Furthermore, SAS-induced accumulation of NO (**Fig. 8D**), ROS (**Fig. 8E**) and lipid-ROS (**Fig. 8F**) was diminished by PDI knockdown for 48 h.

**Figure 8.**
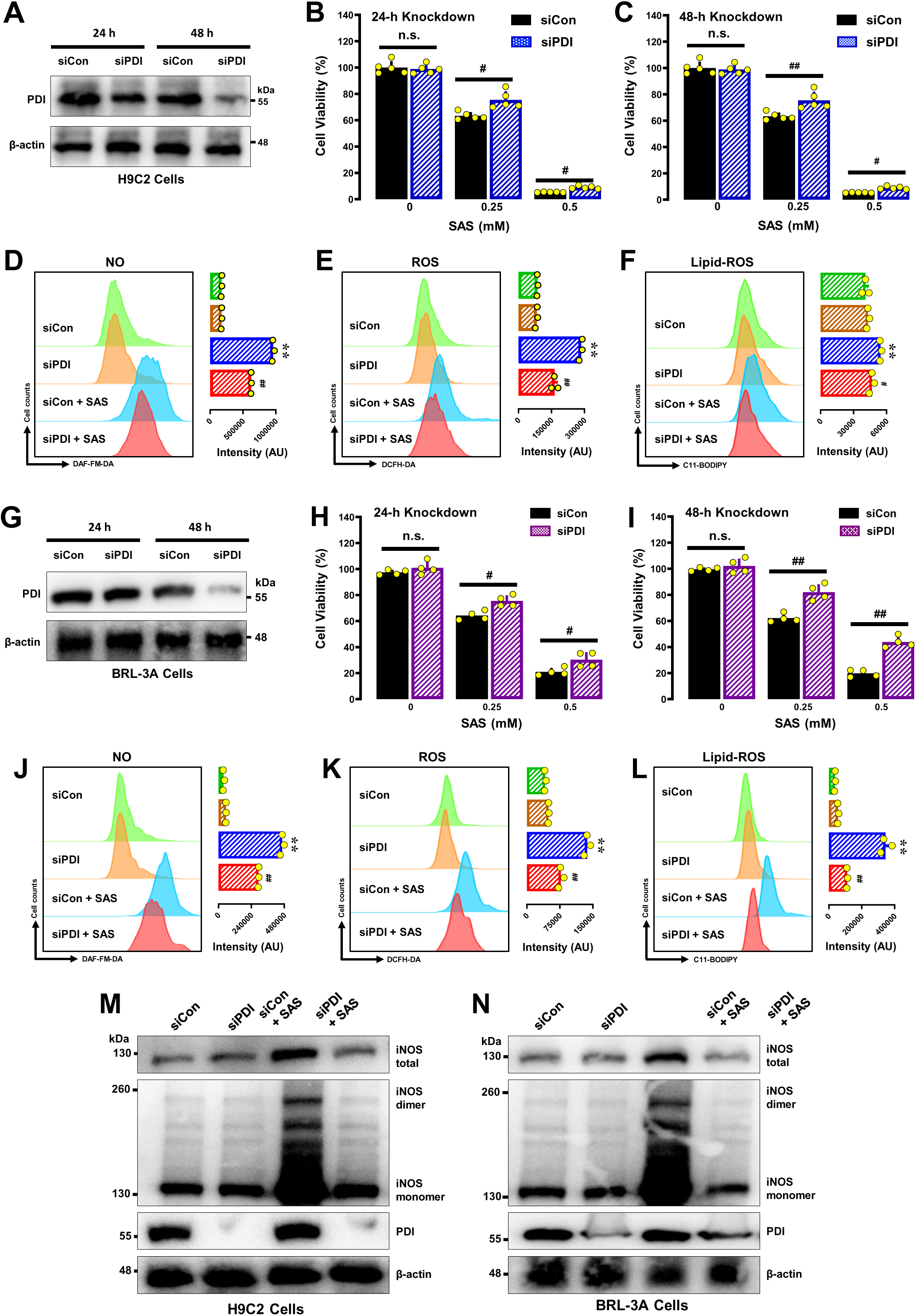
Role of PDI in SAS-induced ferroptosis and accumulation of NO, ROS and lipid-ROS in H9C2 and BRL-3A cells. **A, D.** Effectiveness of PDI-siRNAs in reducing cellular PDI protein level. Cells were transfected with PDI-siRNAs for 24 and 48 h in H9C2 (**A**) and BRL-3A (**D**) cells, respectively, and the PDI protein levels were determined by Western blotting. **B, C, H, I.** Protective effect of PDI knockdown on SAS-induced cytotoxicity in H9C2 (**B, C**) and BRL-3A (**H, I**) cells. Cells were transfected with PDI-siRNAs for 48 h (**C, I**) prior to treatment of cells with SAS (0.5 mM) for an additional 24 h (**B, H**), and cell viability was determined by MTT assay (n = 5). **D, E, F, J, K, L.** Effect of PDI knockdown on SAS-induced NO (**D, J**), ROS (**E, K**) and lipid-ROS (**F, L**) accumulation in H9C2 and BRL-3A cells. Cells were transfected with siPDIs for 48 h prior to treatment with SAS (0.5 mM), and cellular levels of ROS after 8-h SAS exposure were assessed by flow cytometry (the corresponding intensity values are shown on the right; n = 3). **M, N.** Effect of PDI knockdown on total iNOS, its monomer and dimer levels in SAS-treated H9C2 (**M**) and BRL-3A (**N**) cells. Cells were transfected with siCon or PDI siRNAs for 48 h and then treated with 0.5 mM SAS for an additional 6 h. Quantitative data are presented as mean ±SD. * or ^#^ *P* < 0.05; ** or ^##^ *P* < 0.01; n.s., not significant.

Similar observations on the role of PDI in mediating SAS-induced cell death was also made in BRL-3A cells. We found that PDI knockdown (verified by Western blotting, **Fig. 8G**) caused significant reductions in the sensitivity of BRL-3A cells to SAS-induced cytotoxicity (**Fig. 8H** for 24 h; **Fig. 8I** for 48 h). Furthermore, SAS-induced accumulation of NO (**Fig. 8J**), ROS (**Fig. 8K**) and lipid-ROS (**Fig. 8L**) was partially reduced by PDI knockdown.

To investigate whether PDI catalyzes the dimerization of iNOS in SAS-treated cells, we knockdown. We found that PDI knockdown abrogated SAS-induced iNOS dimer levels in these cells (**Fig. 8M** for H9C2 cells; **Fig. 8N** for BRL-3A cells). Notably, PDI knockdown also abrogated SAS-induced upregulation of iNOS total protein levels (**Fig. 8M, 8N**). To provide additional support for the notion that PDI is involved in mediating SAS-induced cell death in both H9C2 and BRL-3A cells, we also examined the protective effects of several known PDI inhibitors, including SNAP, cystamine, 4-OH-E_1_ and BAZ. The results are summarized below.

#### SNAP

SNAP is a *S*-nitrosylating agent which can inhibit the function of PDI by increasing its *S*-nitrosylation (49). In this study, we found that SNAP elicits a modest protection against SAS-induced ferroptosis in H9C2 cells when present at 100 μM (**Fig. 9A, 9B**). SNAP also abrogated SAS-induced accumulation of NO (**Fig. 9C, 9D**), ROS (**Fig. 9E, 9F**) and lipid-ROS (**Fig. 9G, 9H**). In BRL-3A cells, SNAP exhibited a similar protective effect against SAS-induced ferroptosis (**Supplementary Fig. S7A, S7B**), and eliminated the accumulation of NO (**Supplementary Fig. S7C**), ROS (**Supplementary Fig. S7D**) and lipid-ROS (**Supplementary Fig. S7E, S7F**). As expected, SNAP also abrogated SAS-induced increase in iNOS dimer as well as total iNOS protein (**Fig. 9I** for H9C2 cells; **Fig. 9J** for BRL-3A cells).

**Figure 9.**
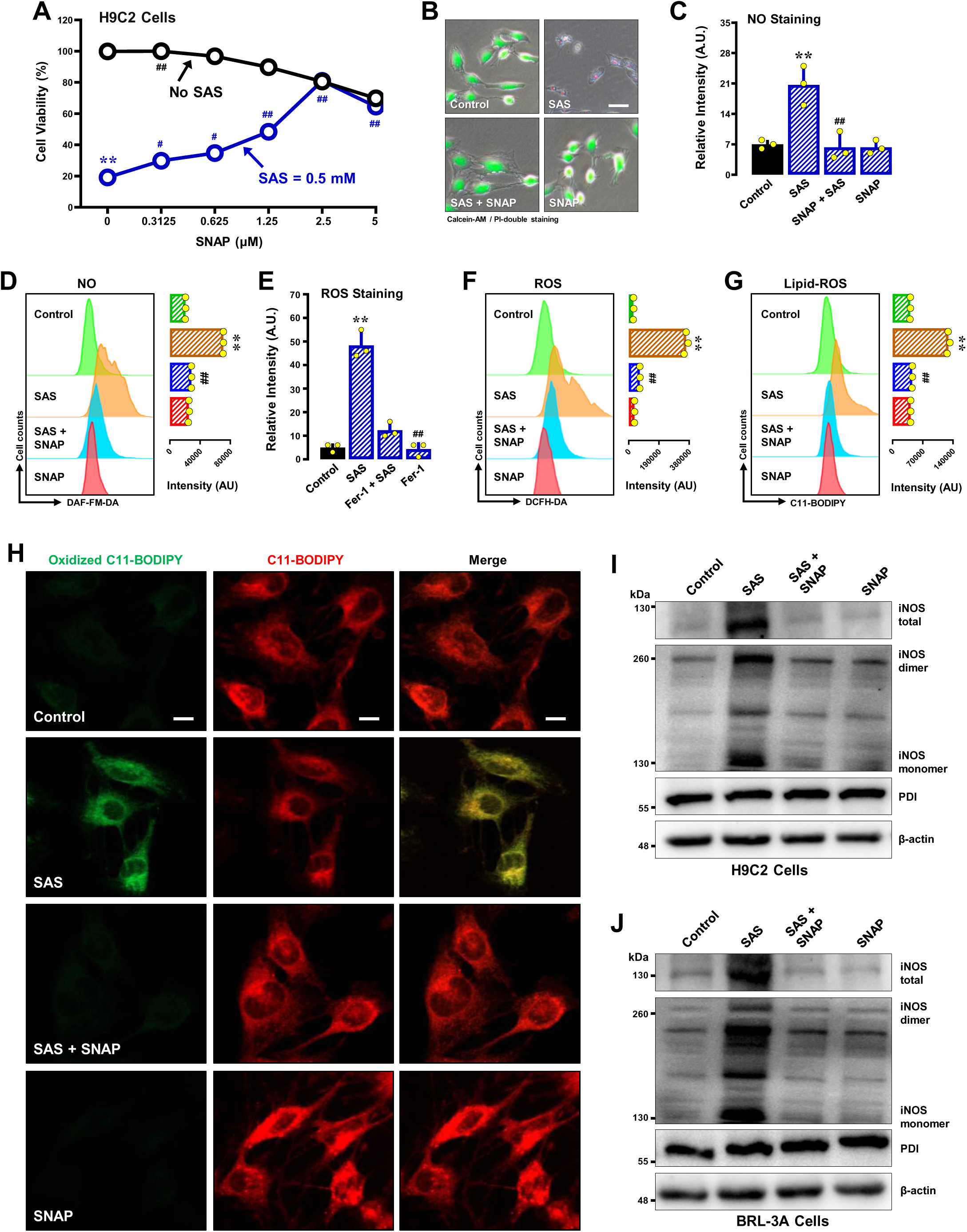
Effect of SNAP on SAS-induced ferroptosis and accumulation of NO, ROS and lipid-ROS in H9C2 and BRL-3A cells. **A, B**. Protective effect of SNAP against SAS-induced cytotoxicity in H9C2 cells. In **A**, cells were treated with SAS (0.5 mM) ±SNAP (0.15625, 0.3125, 0.625, 1.25, 2.5 and 5 µM) for 24 h, and cell viability was determined by MTT assay (n = 5). In **B**, cells were treated with SAS (0.5 mM) ±SNAP (2.5 µM) for 24 h, and then fluorescent images of Calcein-AM/PI-stained cells were captured (green for live cells, and red for dead cells; scale bar = 100 μm). **C, D, E, F, G.** Abrogation by BAZ of SAS-induced accumulation of cellular NO (**C, D**), ROS (**E, F**) and lipid-ROS (**G**) in H9C2 cells. Cells were treated with SAS (0.5 mM) ± SNAP (2.5 µM) for 8 h, and then subjected to flow cytometry (**D, F, G**) and fluorescence microscopy (**C, E**; scale bar = 100 μm). For the fluorescence microscopy data in **C, E**, only the quantitative intensity values are shown (n = 3). For flow cytometry data (**D, F, G**), the corresponding intensity values are shown on the right (n = 3). **H**. Abrogation by SNAP of SAS-induced accumulation of cellular lipid-ROS (confocal microscopy, scale bar = 100 μm) in H9C2 cells. **I, J.** Effect of SNAP on SAS-induced changes in total, monomer and dimer iNOS protein levels in H9C2 (**I**) and BRL-3A (**J**) cells. Cells were treated with SAS (0.5 mM) ±SNAP (2.5 µM) for 8 h, and then levels of the total, monomer and dimer iNOS proteins and PDI were determined by Western blotting. β-Actin was used as a loading control. Quantitative data are presented as mean ±SD. * or ^#^ *P* < 0.05; ** or ^##^ *P* < 0.01; n.s., not significant.

#### Cystamine

Cystamine is a PDI inhibitor that can covalently modify the cysteine residue(s) in PDI’s catalytic region (50), thereby inhibiting its enzymatic activity (23, 25, 51). We found that joint treatment of H9C2 cells with cystamine effectively prevented SAS-induced ferroptotic cell death when cystamine was present at ≥20 μM concentrations (**Fig. 10A** for MTT assay; **Fig. 10B** for Calcein-AM/PI double staining). Cystamine also effectively abrogated SAS-induced accumulation of cellular NO (**Fig. 10C, 10D**), ROS (**Fig. 10E, 10F**) and lipid-ROS (**Fig. 10G, 10H**). Similarly, cystamine prevented SAS-induced ferroptotic cell death in BRL-3A cells (**Supplementary Fig. S8A, S8B**) and reduced the accumulation of NO (**Supplementary Fig. S8C**), ROS (**Supplementary Fig. S8D**) and lipid-ROS (**Supplementary Fig. S8E, S8F**). In addition, cystamine abrogated SAS-induced increase in total iNOS protein levels and their respective dimers (**Fig. 10I** for H9C2 cells; **Fig. 10J** for BRL-3A cells).

**Figure 10.**
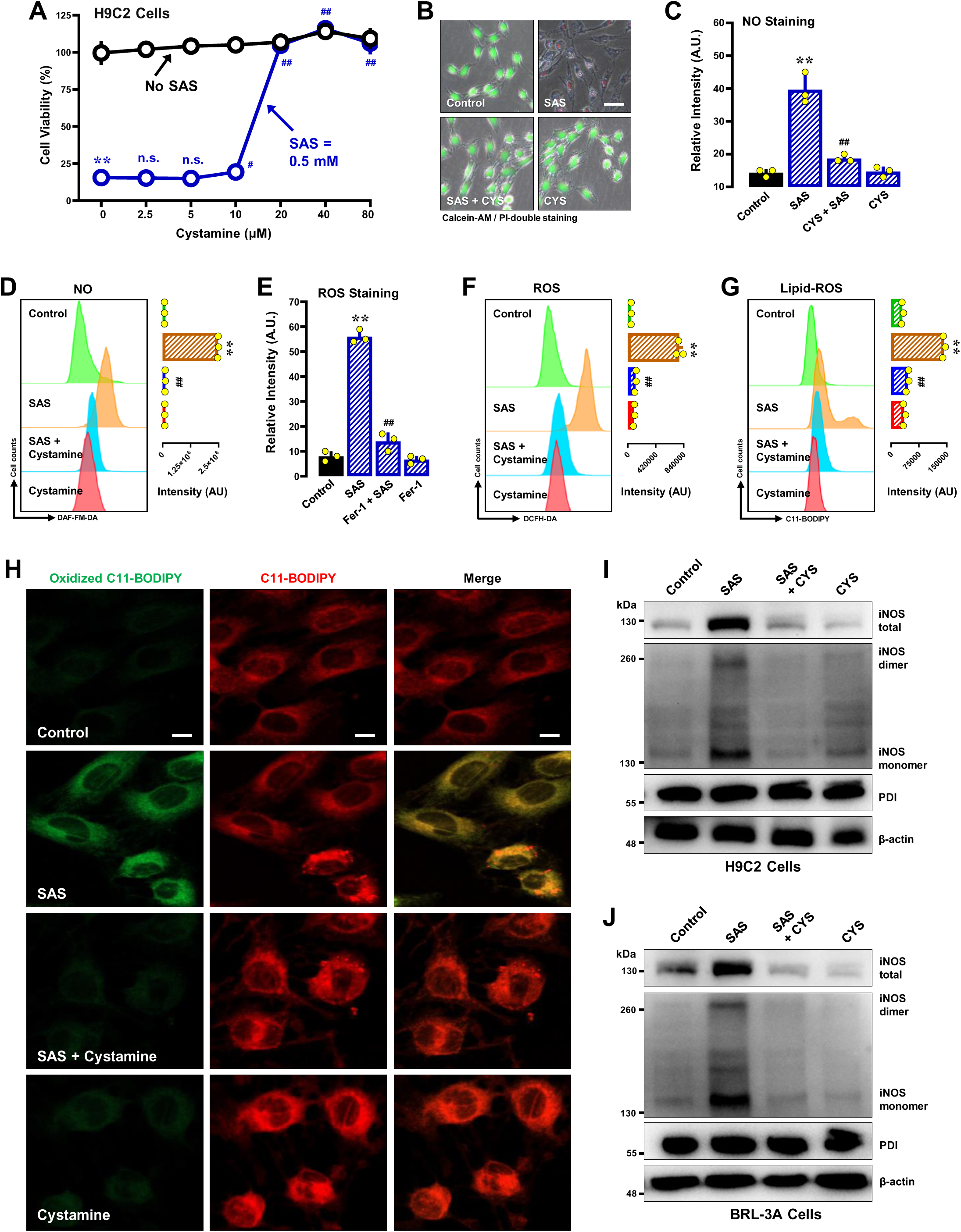
Effect of cystamine on SAS-induced ferroptosis and accumulation of NO, ROS and lipid-ROS in H9C2 and BRL-3A cells. **A, B**. Protective effect of cystamine against SAS-induced cytotoxicity in H9C2 cells. In **A**, cells were treated with SAS (0.5 mM) ±cystamine (2.5, 5, 10, 20, 40 and 80 µM) for 24 h, and then subjected to MTT assay (n = 5). In **B**, cells were treated with SAS (0.5 mM) ± cystamine (40 µM) for 24 h, and then fluorescence images of Calcein-AM/PI-stained cells were captured (green for live cells, and red for dead cells; scale bar = 100 μm). **C, D, E, F, G.** Abrogation by cystamine of SAS-induced accumulation of cellular NO (**C, D**), ROS (**E, F**) and lipid-ROS (**G**) in H9C2 cells. H9C2 cells were treated with SAS (0.5 mM) ± cystamine (40 µM) for 8 h, and then subjected to flow cytometry (**D, F, G**) and fluorescence microscopy (**C, E**; scale bar = 100 μm). For the fluorescence microscopy data in **C, E**, only the quantitative intensity values are shown (n = 3). For flow cytometry data (**D, F, G**), the corresponding intensity values are shown on the right (n = 3). **H**. Abrogation by cystamine of SAS-induced accumulation of cellular lipid-ROS (confocal microscopy, scale bar = 100 μm) in H9C2 cells. **I, J.** Effect of cystamine on SAS-induced changes in total, monomer and dimer iNOS protein levels in H9C2 (**I**) and BRL-3A (**J**) cells. Cells were treated with SAS (0.5 mM) ±cystamine (40 µM) for 8 h, and then levels of the cellular total, monomer and dimer iNOS proteins and PDI were determined by Western blotting. β-Actin was used as a loading control. Quantitative data are presented as mean ±SD. ** or ^##^ *P* < 0.01; n.s., not significant.

#### BAZ

BAZ, a synthetic selective estrogen receptor modulator (52), was recently found to be a noncovalent inhibitor of PDI with a very high potency and can strongly protect cells against chemically-induced ferroptosis (53). We found that BAZ exerts a strong protection against SAS-induced cytotoxicity in H9C2 cells when present at very low concentrations (at ≥156 nM) (**Fig. 12A, 12B**). BAZ effectively abrogated SAS-induced accumulation of cellular NO (**Fig. 12C, 12D**), ROS (**Fig. 12E, 12F**) and lipid-ROS (**Fig. 12G, 12H**). Similar observations were made in BRL-3A cells. BAZ protected SAS-induced ferroptosis at low concentrations (**Supplementary Fig. S9A, S9B**), along with abrogation of SAS-induced accumulation of cellular NO (**Supplementary Fig. S9C**), ROS (**Supplementary Fig. S9D**) and lipid-ROS (**Supplementary Fig. S9E, S9F**). Consistent with its strong ability to inhibit PDI, BAZ was shown to strongly abrogate SAS-induced increase in iNOS dimer levels, along with a reduction in total iNOS protein levels (**Fig. 12I** for H9C2 cells; **Fig. 12J** for BRL-3A cells).

**Figure 11.**
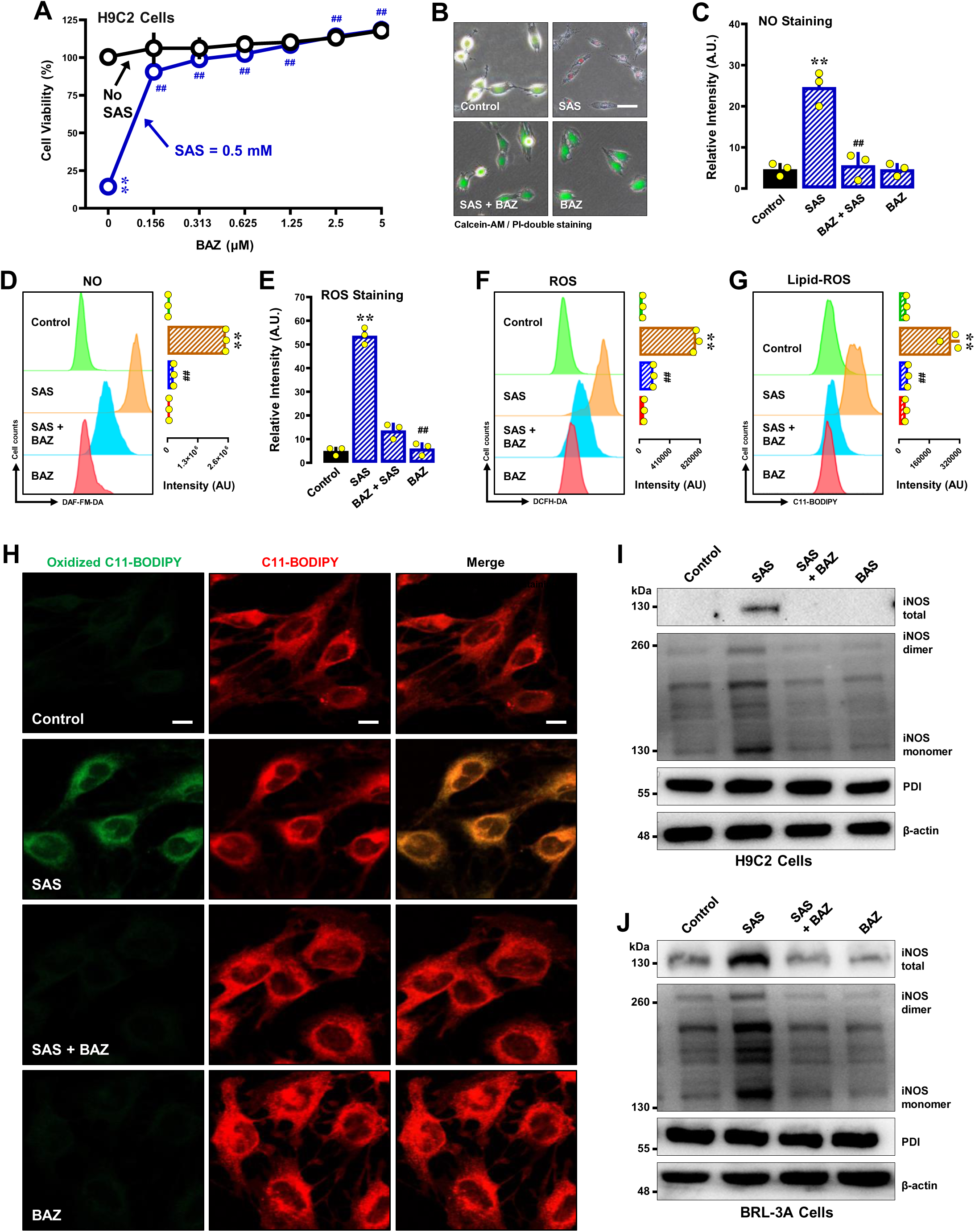
Effect of BAZ on SAS-induced ferroptosis and accumulation of NO, ROS and lipid-ROS in H9C2 and BRL-3A cells. **A, B**. Protective effect of BAZ against SAS-induced cytotoxicity in H9C2 cells. In **A**, cells were treated with SAS (0.5 mM) ±BAZ (0.15625, 0.3125, 0.625, 1.25, 2.5 and 5 µM) for 24 h, and cell viability was determined by MTT assay (n = 5). In **B**, cells were treated with SAS (0.5 mM) ±BAZ (2.5 µM) for 24 h, and then fluorescent images of Calcein-AM/PI-stained cells were captured (green for live cells, and red for dead cells; scale bar = 100 μm). **C, D, E, F, G.** Abrogation by BAZ of SAS-induced accumulation of cellular NO (**C, D**), ROS (**E, F**) and lipid-ROS (**G**) in H9C2 cells. Cells were treated with SAS (0.5 mM) ±BAZ (2.5 µM) for 8 h, and then subjected to flow cytometry (**D, F, G**) and fluorescence microscopy (**C, E**; scale bar = 100 μm). For the fluorescence microscopy data in **C, E**, only the quantitative intensity values are shown (n = 3). For flow cytometry data (**D, F, G**), the corresponding intensity values are shown on the right (n = 3). **H**. Abrogation by BAZ of SAS-induced accumulation of cellular lipid-ROS (confocal microscopy, scale bar = 100 μm) in H9C2 cells. **I, J.** Effect of BAZ on SAS-induced changes in total, monomer and dimer iNOS protein levels in H9C2 (**I**) and BRL-3A (**J**) cells. Cells were treated with SAS (0.5 mM) ±BAZ (2.5 µM) for 8 h, and then levels of the total, monomer and dimer iNOS proteins and PDI were determined by Western blotting. β-Actin was used as a loading control. Quantitative data are presented as mean ±SD. ** or ^##^ *P* < 0.01; n.s., not significant.

**Figure 12.**
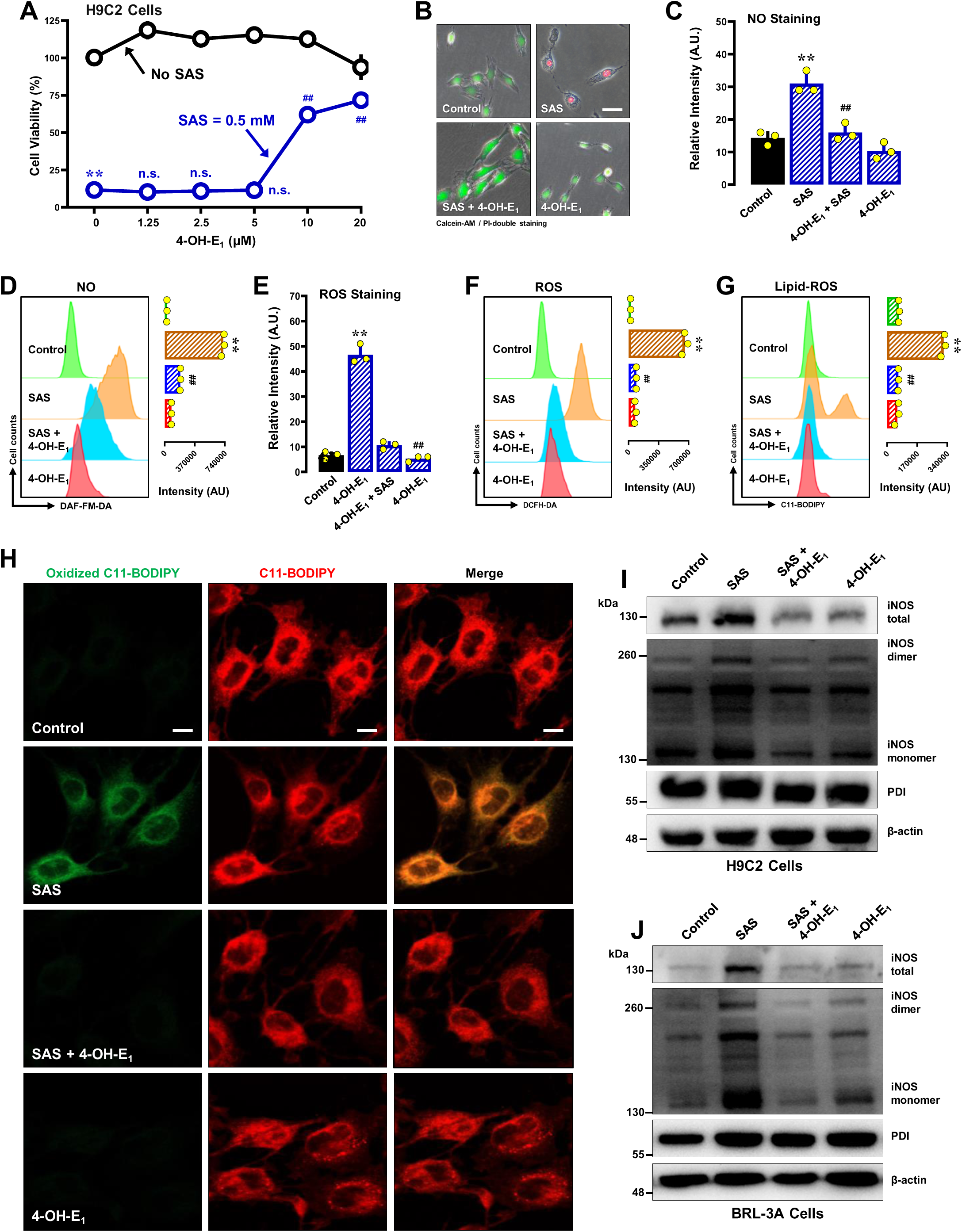
Effect of 4-OH-E_1_ on SAS-induced ferroptosis and accumulation of NO, ROS and lipid-ROS in H9C2 and BRL-3A cells. **A, B**. The protective effect of 4-OH-E_1_ against SAS-induced toxicity was investigated in H9C2 cells. In **A**, cells were exposed to SAS (0.5 mM) ±4-OH-E_1_ (1.25, 2.5, 5, 10 and 20 µM) for a duration of 24 h, and the subsequent assessment of cell viability was conducted through the implementation of the MTT assay (n = 5). In **B**, cells were treated with SAS (0.5 mM) ±4-OH-E_1_ (20 µM) for 24 h, and then fluorescent images of Calcein-AM/PI-stained cells were captured (green for live cells, and red for dead cells; scale bar = 100 μm). **C, D, E, F, G.** Abrogation by 4-OH-E_1_ of SAS-induced accumulation of cellular NO (**C, D**), ROS (**E, F**) and lipid-ROS (**G**) in H9C2 cells. Cells were treated with SAS (0.5 mM) ±4-OH-E_1_ (20 µM) for 8 h, and then subjected to flow cytometry (**D, F, G**) and fluorescence microscopy (**C, E**; scale bar = 100 μm). For the fluorescence microscopy data in **C, E**, only the quantitative intensity values are shown (n = 3). For flow cytometry data (**D, F, G**), the corresponding intensity values are shown on the right (n = 3). **H**. Abrogation by 4-OH-E_1_ of SAS-induced accumulation of cellular lipid-ROS (confocal microscopy, scale bar = 100 μm) in H9C2 cells. **I, J.** Effect of 4-OH-E_1_ on SAS-induced changes in total, monomer and dimer iNOS protein levels in H9C2 (**I**) and BRL-3A (**J**) cells. Cells were treated with SAS (0.5 mM) ±4-OH-E_1_ (20 µM) for 8 h, and then levels of the total, monomer and dimer iNOS proteins and PDI were determined by Western blotting. β-Actin was used as a loading control. Quantitative data are presented as mean ±SD. ** or ^##^ *P* < 0.01; n.s., not significant.

#### 4-OH-E_1_

4-OH-E_1_ is an endogenous estrone metabolite that can non-covalently inhibit DPI’s catalytic function (54). Recently, it was shown that 4-OH-E_1_ can provide a strong protection against chemically-induced ferroptosis (48). In this study, 4-OH-E_1_ was found to have a modest protection against SAS-induced cytotoxicity in H9C2 cells when present at 10–20 μM (**Fig. 11A, 11B**). In a parallel manner, 4-OH-E_1_ also reduced SAS-induced accumulation of NO (**Fig. 11C, 11D**), ROS (**Fig. 11E, 11F**) and lipid-ROS (**Fig. 11G, 11H**). Similar observations were made in BRL-3A cells. We found that 4-OH-E_1_ partially prevented SAS-induced cytotoxicity in BRL-3A cells (**Supplementary Fig. S10A, S10B**). 4-OH-E_1_ also reduced the accumulation of NO (**Supplementary Fig. S10C**), ROS (**Supplementary Fig. S10D**) and lipid-ROS (**Supplementary Fig. S10E, S10F**). In addition, joint treatment of cells with 4-OH-E_1_ also abrogated SAS-induced increases in total iNOS protein levels and their respective dimers (**Fig. 11I** for H9C2 cells; **Fig. 11J** for BRL-3A cells).

Collectively, the above observations with PDI inhibitors underscore the pivotal role of PDI in mediating SAS-induced activation of NOS, the subsequent buildup of cellular NO and ROS/lipid-ROS, and ultimately ferroptotic cell death.

## DISCUSSION

In the past several years, a number of studies have investigated the ferroptosis-inducing ability of SAS in different cell lines in culture, mostly cancer cells (10–12). Recently, we have reported the involvement of PDI in SAS-induced ferroptosis in the immortalized HT22 mouse hippocampal neurons (15). The present study seeks to further characterize the detailed cellular and biochemical mechanisms of SAS-induced ferroptotic cell death in two additional cell lines, namely, the H9C2 cardiomyocytes and BRL-3A hepatocytes. Summarized below is the experimental evidence presented in this study which jointly offers support for the notion that the PDI–NOS–NO–ROS/lipid–ROS pathway plays a pivotal role in mediating SAS-induced oxidative cytotoxicity.

### SAS induces sequential NO, ROS and lipid-ROS accumulation

First, it is confirmed in this study that SAS-induced cell death closely resembles oxidative ferroptosis based on the observations that SAS-induced cell death can be effectively rescued by Fer-1 and DFO but not by Nec-1 and z-VAD-FMK. Next, a series of experimental evidence is presented in this study for the hypothesis that NO accumulation in SAS-treated cells is an early event which then results in the accumulation of cellular ROS and lipid-ROS, and ultimately ferroptotic cell death. ***i.*** A modest increase in NO levels is observed in H9C2 and BRL-3A cells at 1–2 h following SAS exposure, a significant buildup of cellular ROS and lipid-ROS is evident at ∼4h. This observation suggests that in SAS-treated cells, cellular NO buildup takes place first, and is followed by accumulation of cellular ROS and lipid-ROS. ***ii.*** SNP, which can directly release NO intracellularly (43), is shown to strongly increase the sensitivity of H9C2 and BRL-3A cells to SAS-induced cytotoxicity. This heightened sensitivity is attributed to an accelerated buildup of cellular ROS and lipid-ROS, consequent to a swift escalation in cellular NO levels. ***iii.*** cPTIO, an NO scavenger (37), not only mitigates SAS-induced NO accumulation but also decreases the accumulation of ROS and lipid-ROS triggered by SAS. The protective effect of cPTIO is associated with its ability to counteract SAS-induced cell death. Collectively, these observations provide substantial evidence for the hypothesis that NO is a more upstream mediator than ROS and lipid-ROS in SAS-induced ferroptois in cultured cells.

It has been established that the accumulation of cellular ROS, particularly cellular lipid-ROS, constitutes a pivotal aspect of chemically-induced oxidative ferroptosis (16, 18, 56). In this study, we demonstrate that the accumulation of ROS and lipid-ROS in SAS-treated H9C2 and BRL-3A cells also plays an important role in ferroptosis progression. For instance, NAC, a well-known antioxidant (39, 40), can effectively eliminate the accumulation of cellular NO, ROS and lipid-ROS in SAS-treated cells, thereby providing a robust cytoprotection. Similarly, Fer-1, an aromatic primary amine with the capacity to suppress lipid peroxidation (28), can also effectively rescue SAS-induced ferroptosis in H9C2 and BRL-3A cells. It is of note that while Fer-1 potently suppresses SAS-induced lipid-ROS accumulation, its effect on SAS-induced NO and ROS accumulation is comparatively modest. This is not unexpected, given that Fer-1 is predominantly a scavenger of cellular lipid-ROS, which is downstream of the cellular NO and ROS in the proposed iNOS→NO→ROS→lipid-ROS cascade. Collectively, this study reveals that SAS-induced cellular NO accumulation occurs prior to cellular ROS and lipid-ROS accumulation, and cellular NO, ROS and lipid-ROS jointly drive oxidative ferroptosis in SAS-treated H9C2 and BRL-3A cells.

### PDI mediates NOS activation in SAS-treated cells

NOS catalyzes the production of NO by oxidizing *L*-arginine to *L*-citrulline (57, 58). Notably, iNOS exists in two structural forms: a dimeric iNOS, which is catalytically active and essential for NO synthesis, and a monomeric iNOS, which is catalytically inactive. In this study, we find that SAS treatment increases iNOS dimerization in a concentration- and time-dependent manner in H9C2 and BRL-3A cells. In addition, we show that treatment of these cells with SAS results in a marked upregulation of the total cellular iNOS protein levels. It is known that upregulation of iNOS and particularly increased iNOS dimerization can result in elevated NO production, which, under certain conditions, can be highly cytotoxic (59, 60). Interestingly, despite these changes in iNOS levels and activity, the cellular levels of PDI, which is the enzyme that catalyzes iNOS dimerization, remains largely unaffected.

To provide experimental support for the suggestion that iNOS dimerization leads to NO and ROS/lipid-ROS accumulation and ultimately cell death, we jointly treat H9C2 and BRL-3A cells with SMT, an iNOS inhibitor, and find that SMT attenuates SAS-induced iNOS upregulation and dimerization as well as NO accumulation, which is accompanied by reductions in SAS-induced ROS/lipid-ROS accumulation and ferroptosis. In a similar fashion, iNOS knockdown also partially eliminates SAS cytotoxicity. Jointly, these results demonstrate that SAS-induced iNOS upregulation and dimerization contribute critically to SAS-induced ferroptosis.

In this study, we have also provided evidence to show that PDI is a pivotal upstream mediator of SAS-induced ferroptosis by catalyzing iNOS dimerization, which then leads to NO and ROS/lipid-ROS accumulation, and ultimately, oxidative cell death. Specifically, we find that knockdown of cellular PDI with specific PDI siRNAs strongly attenuates SAS-induced NO and ROS/lipid-ROS accumulation, along with a protection of SAS-induced cell death.

Similar to the effect of PDI knockdown, we find that several PDI inhibitors also exhibit a protective effect against SAS-induced ferroptosis in H9C2 and BRL-3A cells. SNAP, a thiol-nitrosylating agent (49), can enhance DPI *S*-nitrosylation (49). In our previous work, we showed that PDI predominantly exists in the *S*-nitrosylated state in untreated HT22 cells, and exposure to glutamate induces PDI *S*-denitrosylation (49). This shift promotes PDI oxidation and activation, enabling it to catalyze NOS dimerization. Earlier we reported that SNAP, which has some cytotoxicity when present alone (21, 23, 24), partially rescued glutamate-induced oxidative cytotoxicity (oxytosis) through enhanced PDI *S*-nitrosylation (21). In addition, a similar rescuing effect of SNAP was observed during erastin-induced ferroptosis (23, 24). In the present study, we find that SNAP also provides significant protection against SAS-induced ferroptosis, which is associated with suppression of SAS-induced iNOS upregulation and dimerization. As expected, these effects are associated with a marked decrease in cellular NO, ROS and lipid-ROS levels.

Like SNAP, we find that cystamine, a covalent inhibitor of PDI (25, 61, 62), can strongly attenuate SAS-induced iNOS dimerization and cellular NO, ROS/lipid-ROS accumulation, which is accompanied by ferroptosis protection. Additionally, cystamine also inhibits SAS-induced iNOS upregulation, which further contributes to its cytoprotective properties. It is important to note that cystamine is not a reducing agent (rather it is a mild oxidizing agent), and it can covalently modify the free thiol groups in PDI’s catalytic sites, resulting in inhibition of PDI’s catalytic function. The strong protective effect of cystamine underscores the pivotal upstream role of PDI in SAS-induced ferroptosis.

Different from cystamine, BAZ and 4-OH-E_1_ can inhibit PDI’s catalytic activities through non-covalent binding interactions with PDI (48, 53). BAZ displays a strong protection against SAS-induced cell death. The cytoprotective effect is closely associated with its inhibition of PDI-mediated iNOS dimerization in SA-treated cells, along with abrogation of cellular NO, ROS and lipid-ROS accumulation. Like BAZ, 4-OH-E_1_ also suppresses PDI-mediated iNOS dimerization in SAS-treated cells, which is associated with reduced accumulation of cellular NO, ROS and lipid-ROS. Here, it should be noted that 4-OH-E_1_ has a relatively weaker efficacy compared to BAZ in abrogating SAS-induced ROS buildup and cell death because 4-OH-E_1_ itself is a catechol (which is chemically reactive) and can undergo oxidation/auto-oxidation to generate ROS under certain conditions (63). Together, the experimental evidence discussed here with several PDI inhibitors offers additional support for the pivotal role of PDI in mediating SAS-induced ferroptosis. Additionally, these results also highlight the importance of PDI as a crucial drug target for protection against oxidative ferroptosis.

Lastly, regarding the potential causes of PDI activation (*i.e.*, oxidation) in SAS-treated cells, first it is ruled out in this study that PDI can be directly activated by SAS (**Supplementary Fig. S6**). As depicted in **Supplementary Fig. S11**, it is understood that PDI is favored to be in the oxidized state when cellular free GSH levels are reduced, such as in the presence of SAS or erastin. These chemicals can slowly deplete cellular free GSH, and because of this slow process, PDI activation and the subsequent NOS dimerization also take some time to happen. However, in the case of RSL3, which is an inhibitor of TrxR1 (24, 64), PDI activation and NOS dimerization can occur more quickly, and as a result, RSL3-induced ferroptosis also occurs much faster (24).

### Conclusions

As summarized in **Supplementary Fig. S11**, the present study has shown that SAS is capable of inducing ferroptosis in H9C2 cardiomyocytes and BRL-3A hepatocytes in culture, and this process is associated with time-dependent, sequential increases in cellular NO, ROS and lipid-ROS levels. Treatment of these cells with SAS activates PDI-mediated iNOS dimerization, which activates the catalytic activity of iNOS for NO production. Furthermore, SAS also upregulates iNOS protein levels in these cells, which also contributes to elevated NO production and accumulation. Genetic manipulation of PDI expression and pharmacological inhibition of PDI’s catalytic activity can effectively abrogate SAS-induced iNOS dimerization and cellular NO, ROS and lipid-ROS accumulation, along with strong protection against ferroptotic cell death. Collectively, the findings of this study, along with our recent observations (15), jointly demonstrate a pivotal role of PDI in SAS-induced ferroptosis through the activation of the PDI→NOS→NO→ROS/lipid-ROS pathway, and also offer new strategies to sensitizing cancer cells to SAS-induced ferroptosis, such as through the use of NO-releasing agents or TrxR1 inhibitors.

## DECLARATION OF INTEREST STATEMENT

All authors of this study do not have any conflict of interest to declare.

---

**Supplementary Fig. S1.**
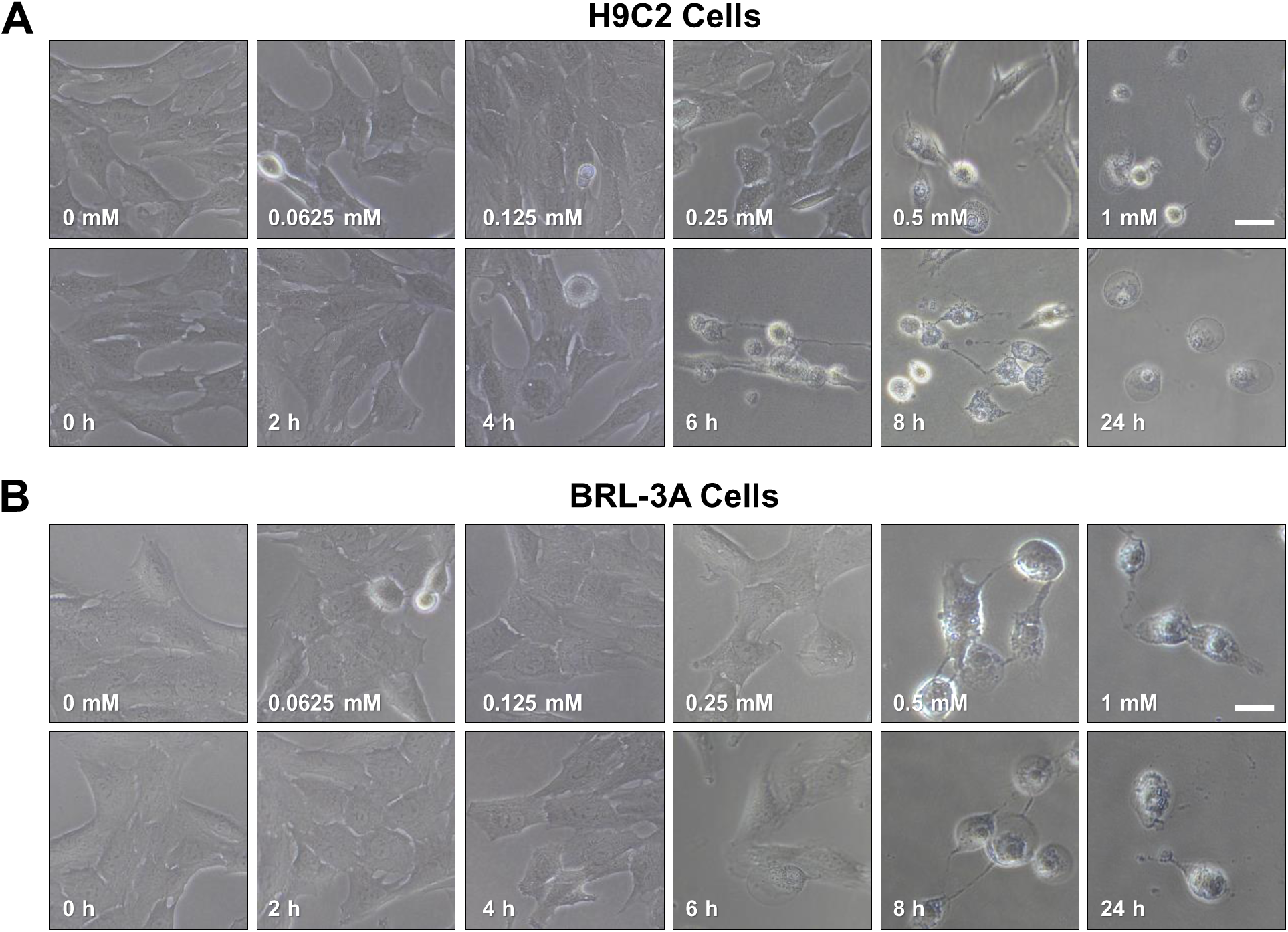
Concentration- and time-dependent induction of ferroptotic cell death by SAS in H9C2 and BRL-3A cells. **A.** Concentration- and time-dependent changes in gross morphology of H9C2 cells after exposure to increasing concentrations of SAS for 24 h (**upper panel**) or 0.5 mM SAS for varying durations (**lower panel**). The cellular images were captured with a light microscope (40×, scale bar = 100 μm). **B.** Concentration- and time-dependent changes in gross morphology of BRL-3A cells after exposure to increasing concentrations of SAS for 24 h (**upper panel**) or 0.5 mM SAS for varying durations (**lower panel**). The cellular images were captured with a light microscope (40×, scale bar = 100 μm).

**Supplementary Fig. S2.**
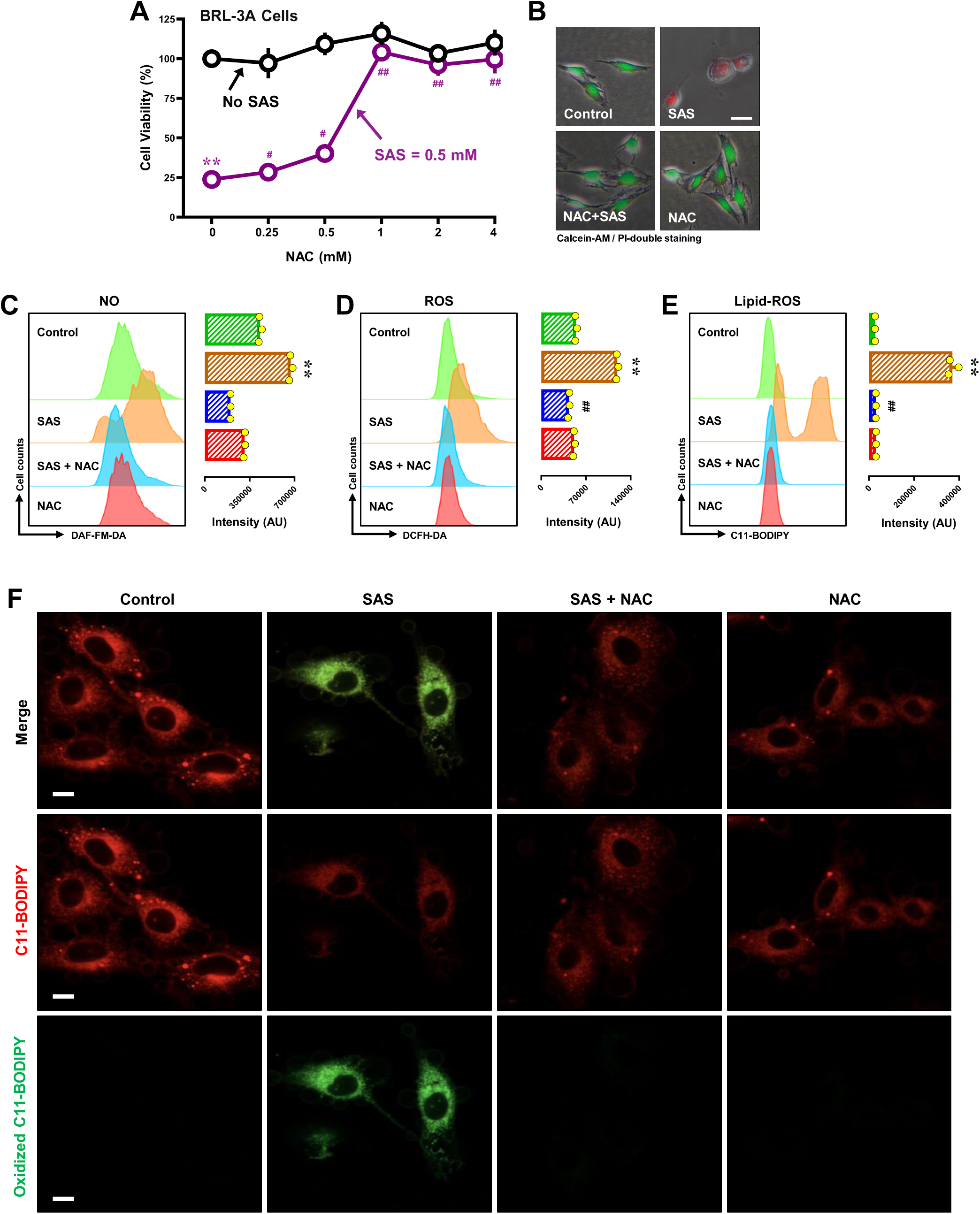
Effect of NAC on SAS-induced ferroptosis and accumulation of NO, ROS and lipid-ROS in BRL-3A cells. **A, B.** Protective effect of NAC against SAS-induced cytotoxicity. In **A**, cells were treated with SAS (0.5 mM) ± NAC (0.125, 0.25, 0.5, 2, 2 and 4 mM) for 24 h, and then subjected to MTT assay (n = 4). In **B**, cells were treated with SAS (0.5 mM) ± NAC (0.5 mM) for 24 h, and then fluorescent images of Calcein-AM/PI-stained cells were captured (green for live cells, and red for dead cells; scale bar = 100 μm). **C, D, E**. Abrogation by NAC of SAS-induced accumulation of cellular NO (**C**), ROS (**D**) and lipid-ROS (**E**). Cells were treated with SAS (0.5 mM) ± NAC (0.5 mM) for 8 h, and then subjected to analytical flow cytometry. The left panels are the histograms, and the right panels are the quantitative values (n = 3). **F**. Abrogation by SNAP of SAS-induced accumulation of cellular lipid-ROS (confocal microscopy, scale bar = 100 μm). Quantitative data are presented as mean ± SD. * or ^#^ *P* < 0.05; ** or ^##^ *P* < 0.01; n.s., not significant.

**Supplementary Fig. S3.**
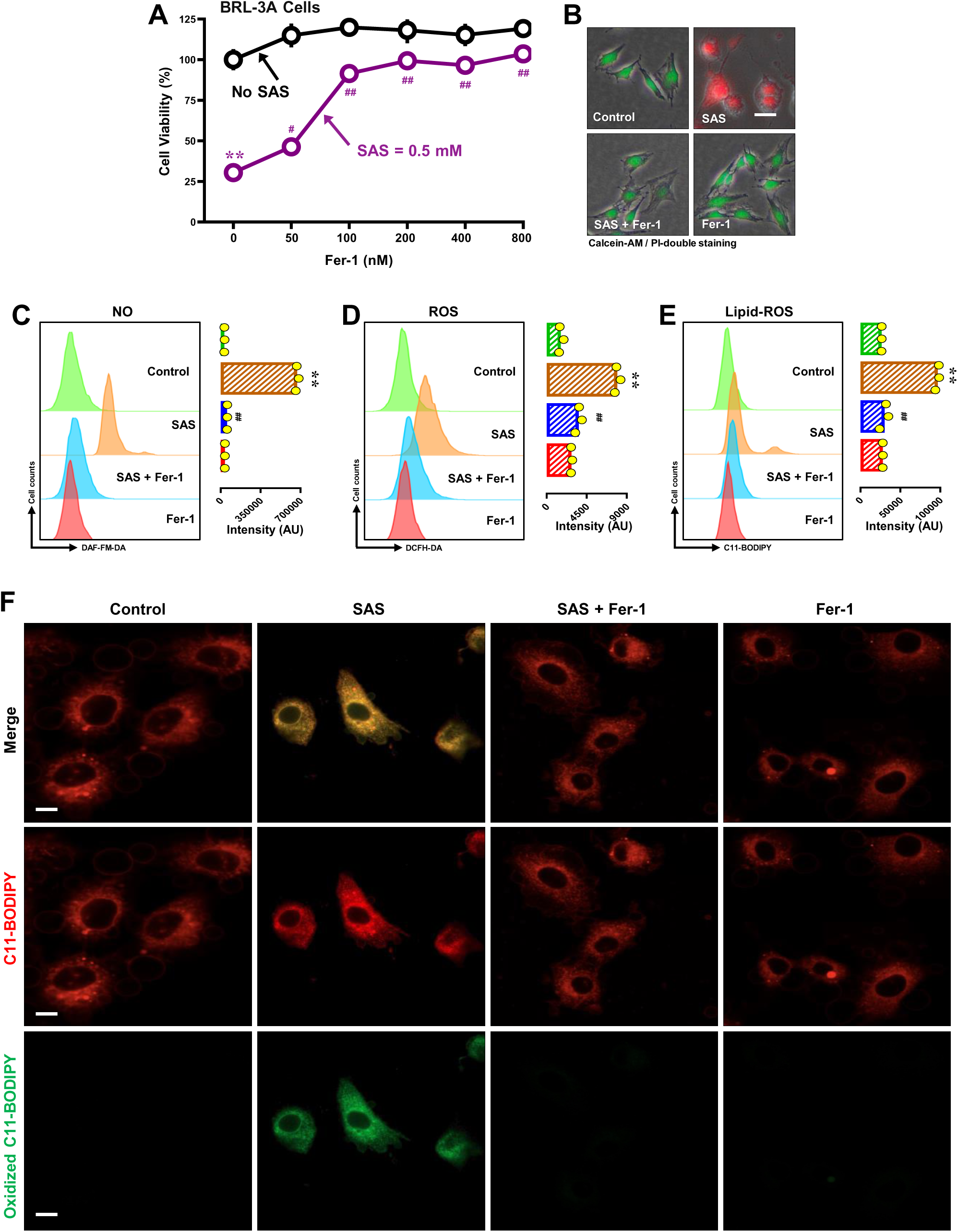
Effect of Fer-1 on SAS-induced ferroptosis and accumulation of NO, ROS and lipid-ROS in BRL-3A cells. **A, B.** Protective effect of Fer-1 against SAS-induced cytotoxicity. In **A**, cells were treated with SAS (0.5 mM) ± Fer-1 (25, 50, 100, 200, 400 and 800 nM) for 24 h, and then cell viability was determined by MTT assay (n = 4). In **B**, cells were treated with SAS (0.5 mM) ± Fer-1 (200 nM) for 24 h, and then fluorescent images of Calcein-AM/PI-stained cells were captured (green for live cells, and red for dead cells; scale bar = 100 μm). **C, D, E**. Abrogation by Fer-1 of SAS-induced accumulation of cellular NO (**C, D**), ROS (**E, F**) and lipid-ROS (**G**). Cells were treated with SAS (0.5 mM) ± Fer-1 (200 nM) for 8 h, and then subjected to analytical flow cytometry. The left panels are the histograms, and the right panels are the quantitative values (n = 3). **F**. Abrogation by SNAP of SAS-induced accumulation of cellular lipid-ROS (confocal microscopy, scale bar = 100 μm). Quantitative data are presented as mean ± SD. * or ^#^ *P* < 0.05; ** or ^##^ *P* < 0.01; n.s., not significant.

**Supplementary Fig. S4.**
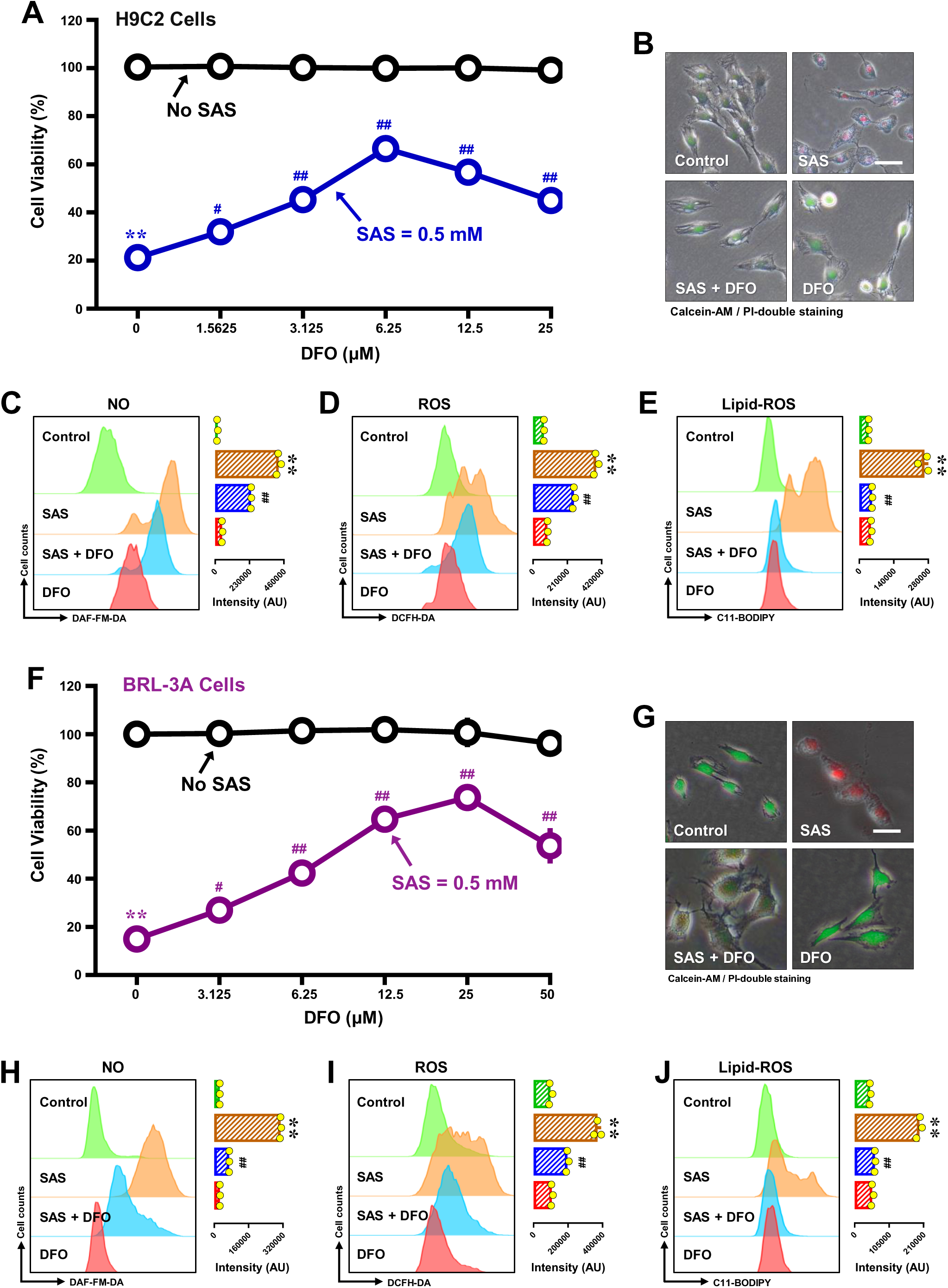
Effect of DFO on SAS-induced ferroptosis and accumulation of NO, ROS and lipid-ROS in H9C2 and BRL-3A cells. **A, B.** Effect of DFO against SAS-induced cytotoxicity in H9C2 cells. In **A**, cells were treated with SAS (0.5 mM) ± DFO (1.563, 3.125, 6.25, 12.5 and 25 μM) for 24 h, and then cell viability was determined by MTT assay (n = 5). In **B**, cells were treated with SAS (0.5 mM) ± DFO (6.25 μM) for 24 h, and then fluorescent images of Calcein-AM/PI-stained cells were captured (green for live cells, and red for dead cells; scale bar = 100 μm). **C, D, E**. Partial abrogation by DFO of SAS-induced accumulation of cellular NO (**C**), ROS (**D**) and lipid-ROS (**E**) in H9C2 cells. Cells were treated with SAS (0.5 mM) ± DFO (6.25 μM) for 8 h, and then subjected to flow cytometry. The left panels are the histograms, and the right panels are the quantitative values (n = 3). **F, G.** Effect of DFO against SAS-induced cytotoxicity in BRL-3A cells. In **F**, cells were treated with SAS (0.5 mM) in the presence or absence of varying concentrations of DFO (3.125, 6.25, 12.5, 25, and 50 μM) for 24 h, and cell viability was assessed using the MTT assay (n = 5). In **G**, cells were treated with SAS (0.5 mM) ± DFO (25 μM) for 24 h, followed by fluorescence imaging of Calcein-AM/PI-stained cells, where green indicates live cells and red indicates dead cells (scale bar = 100 μm). **H, I, J**. DFO suppresses SAS-induced accumulation of cellular NO (**H**), ROS (**I**), and lipid-ROS (**J**) in BRL-3A cells. Cells were treated with SAS (0.5 mM) ± DFO (25 μM) for 8 h, followed by flow cytometry analysis. The left panels are the histograms, and the right panels are the quantitative values (n = 3). Quantitative data are presented as mean ± SD. * or ^#^ *P* < 0.05; ** or ^##^ *P* < 0.01; n.s., not significant.

**Supplementary Fig. S5.**
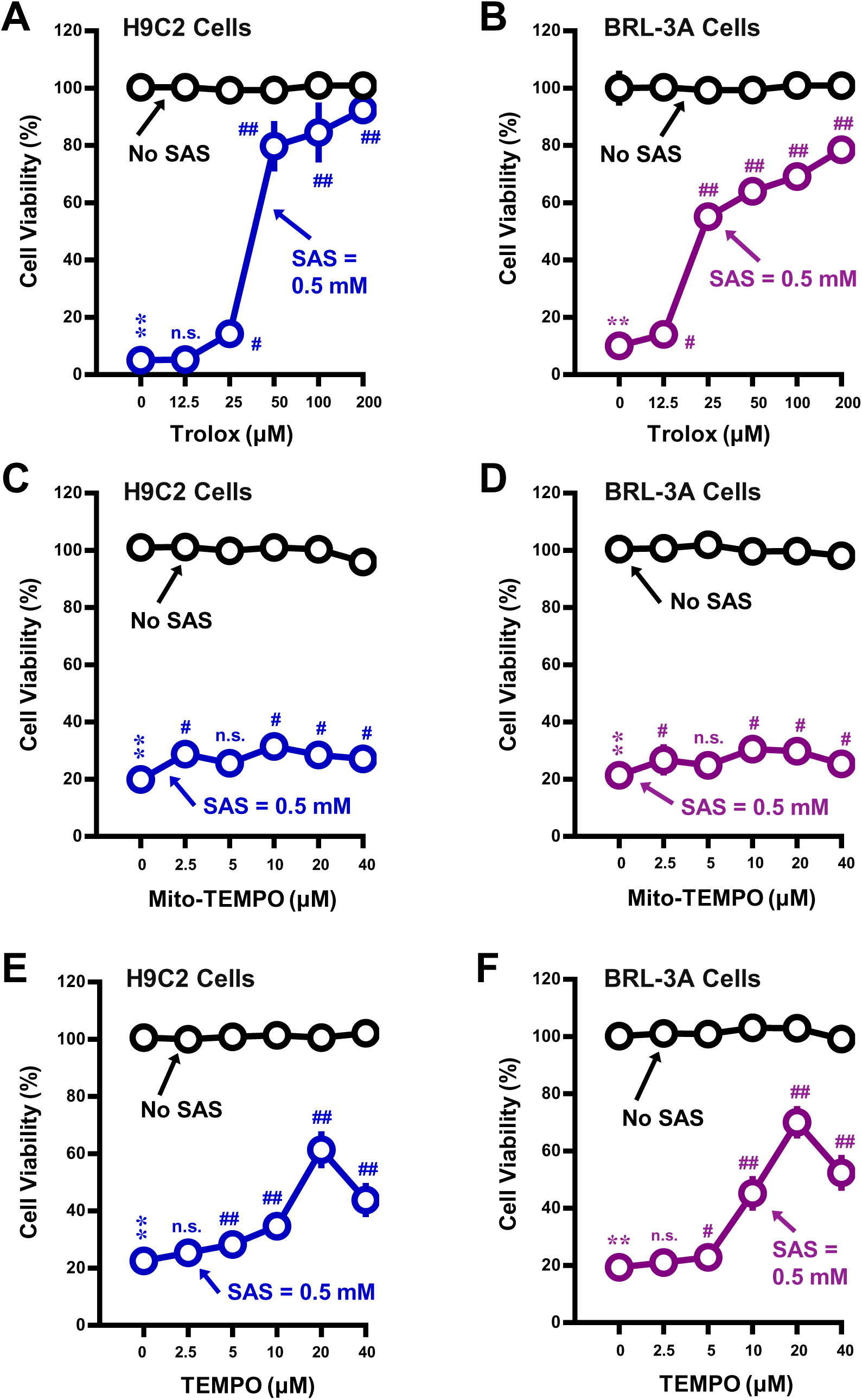
Effect of Trolox, MitoTEMPO and TEMPO on SAS-induced death in H9C2 and BRL-3A cells. **A, B.** Protective effect of Trolox against SAS-induced cytotoxicity following treatment of H9C2 (**A**) and BRL-3A (**B**) cells with SAS (0.5 mM) ± Trolox (12.5, 25, 50, 100 and 200 µM) for 24 h (MTT assay, n = 5). **C, D.** Lack of a protective effect of MitoTEMPO against SAS cytotoxicity. Cells were treated with SAS (0.5 mM) ± MitoTEMPO (2.5, 5 10, 20 and 40 μM) for 24 h, and cell viability was determined by MTT assay (n = 5). **E, F.** Partial protective effect of TEMPO against SAS cytotoxicity. Cells were treated with SAS (0.5 mM) ± TEMPO (2.5, 5 10, 20 and 40 μM) for 24 h, and cell viability was determined by MTT assay (n = 5). Quantitative data are presented as mean ± SD. * or ^#^ *P* < 0.05; ** or ^##^ *P* < 0.01; n.s., not significant.

**Supplementary Fig. S6.**
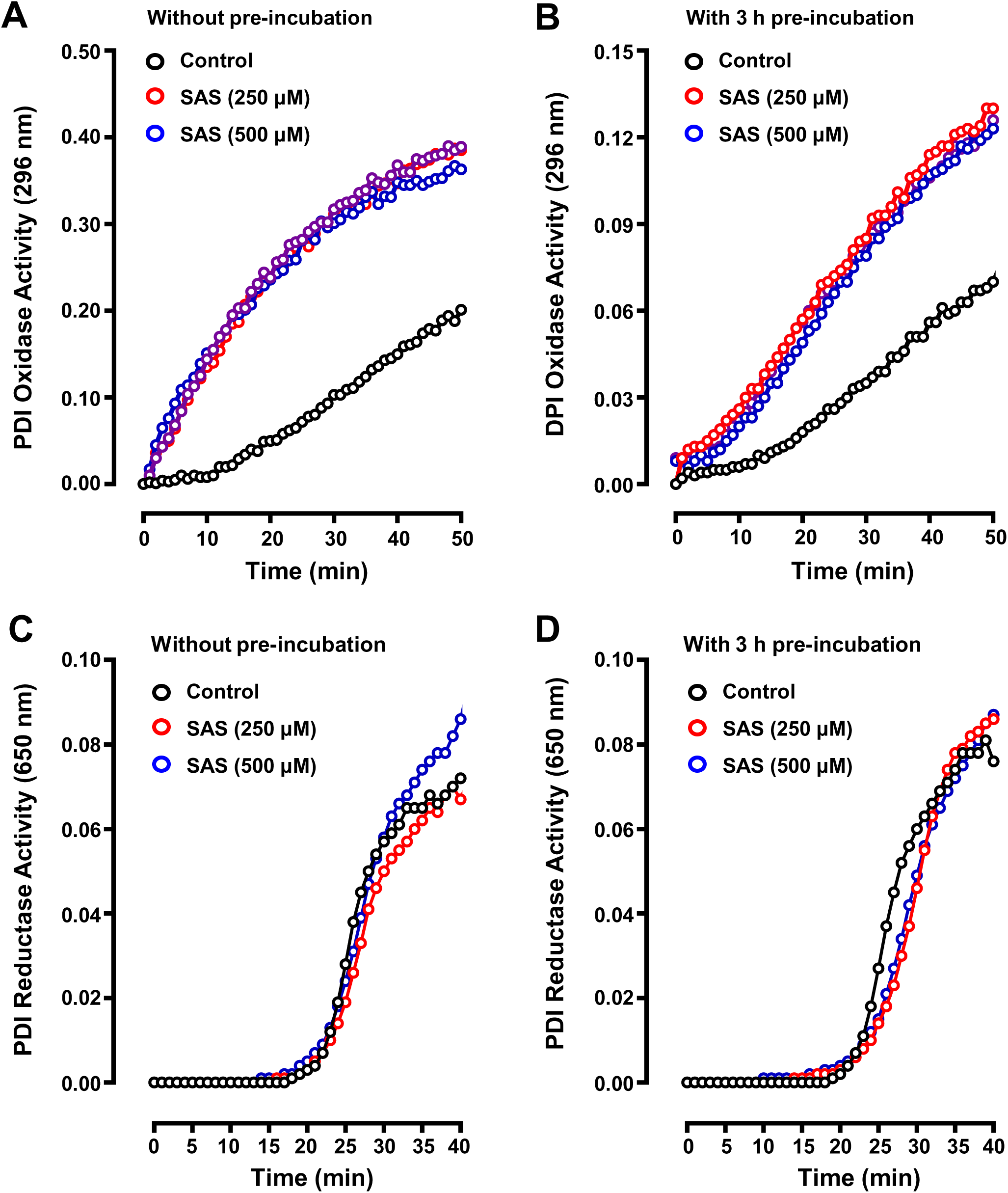
Effect of SAS on the oxidase and reductase activities of PDI in the *in-vitro* enzymatic assays. **A, B.** Effect of SAS (at 250 and 500 μM) on the oxidase activity of PDI (**A, B**). The oxidase activity of PDI was assayed by measuring PDI-mediated RNase A refolding *in vitro*. The assay was repeated to confirm the observations. The data from one representative assay is shown. **A, B.** Effect of SAS (at 250 and 500 μM) of the reductase activity of PDI. The reductase activity of PDI was assayed by measuring PDI-mediated insulin aggregation *in vitro*. The assay was repeated to confirm the observations. The data from one representative assay is shown.

**Supplementary Fig. S7.**
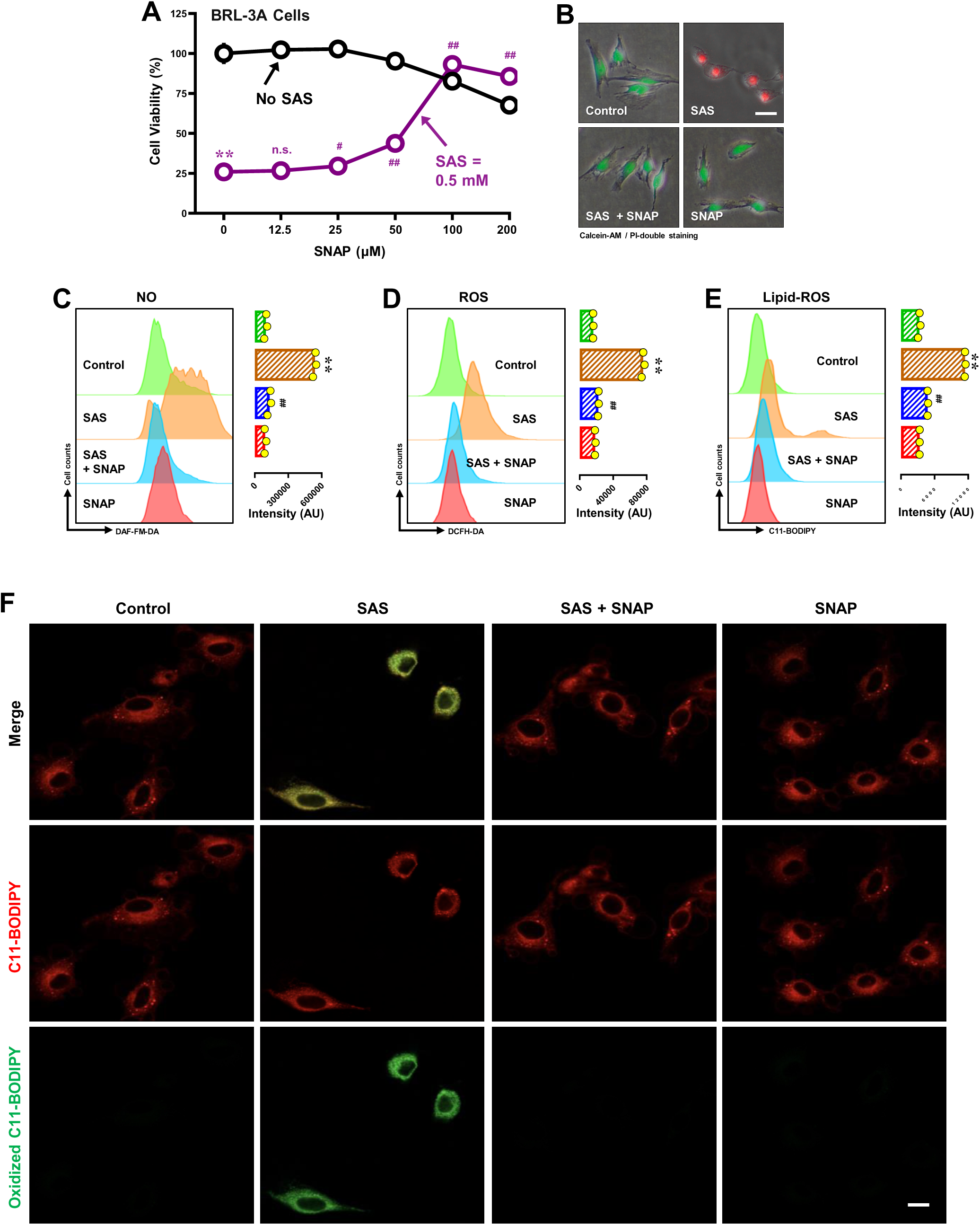
Effect of SNAP on SAS-induced ferroptosis and accumulation of NO, ROS and lipid-ROS in BRL-3A cells. **A, B**. Protective effect of SNAP against SAS-induced cytotoxicity. In **A**, cells were treated with SAS (0.5 mM) ± SNAP (12.5, 25, 50, 100 and 200 µM) for 24 h, and cell viability was determined by MTT assay (n = 5). In **B**, cells were treated with SAS (0.5 mM) ± SNAP (100 µM) for 24 h, and then fluorescent images of Calcein-AM/PI-stained cells were captured (green for live cells, and red for dead cells; scale bar = 100 μm). **C, D, E.** Abrogation by SNAP of SAS-induced accumulation of cellular NO (**C**), ROS (**D**) and lipid-ROS (**E**). Cells were treated with SAS (0.5 mM) ± SNAP (100 µM) for 8 h, and then subjected to analytical flow cytometry. The left panels are the histograms, and the right panels are the quantitative values (n = 3). **F**. Abrogation by SNAP of SAS-induced accumulation of cellular lipid-ROS (confocal microscopy, scale bar = 100 μm). Quantitative data are presented as mean ± SD. * or ^#^ *P* < 0.05; ** or ^##^ *P* < 0.01; n.s., not significant.

**Supplementary Fig. S8.**
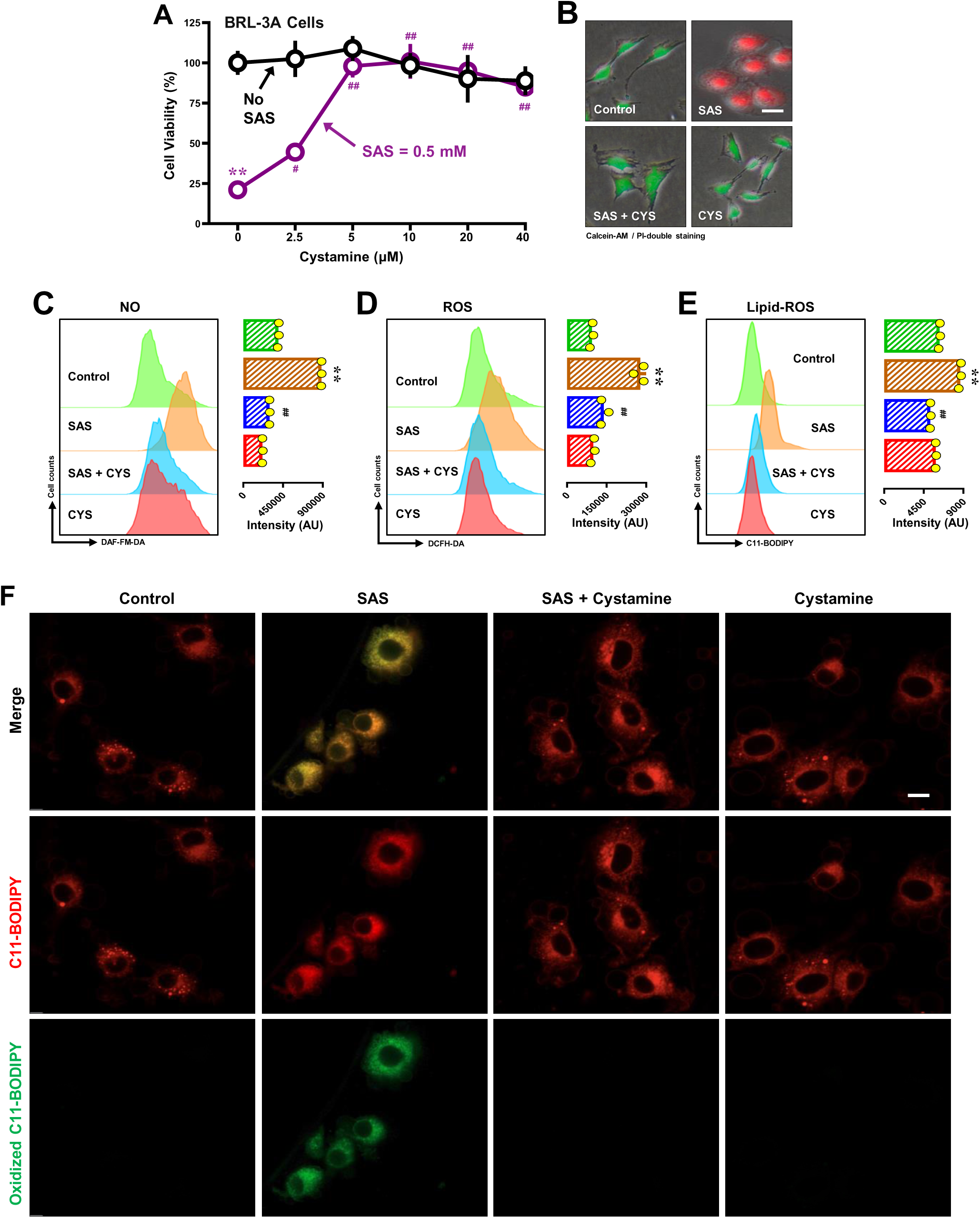
Effect of cystamine on SAS-induced ferroptosis and accumulation of NO, ROS and lipid-ROS in BRL-3A cells. **A, B**. Protective effect of cystamine against SAS-induced cytotoxicity. In **A**, cells were treated with SAS (0.5 mM) ± cystamine (2.5, 5, 10, 20 and 40 µM) for 24 h, and cell viability was determined by MTT assay (n = 5). In **B**, cells were treated with SAS (0.5 mM) ± cystamine (10 µM) for 24 h, and then fluorescence images of Calcein-AM/PI-stained cells were captured (green for live cells, and red for dead cells; scale bar = 100 μm). **C, D, E.** Abrogation by cystamine of SAS-induced accumulation of cellular NO (**C,**), ROS (**D**) and lipid-ROS (**E**). Cells were treated with SAS (0.5 mM) ± cystamine (10 µM) for 8 h, and then subjected to analytical flow cytometry. The left panels are the histograms, and the right panels are the quantitative values (n = 3). **F**. Abrogation by cystamine of SAS-induced accumulation of cellular lipid-ROS (confocal microscopy, scale bar = 100 μm). Quantitative data are presented as mean ± SD. ** or ^##^ *P* < 0.01; n.s., not significant.

**Supplementary Fig. S9.**
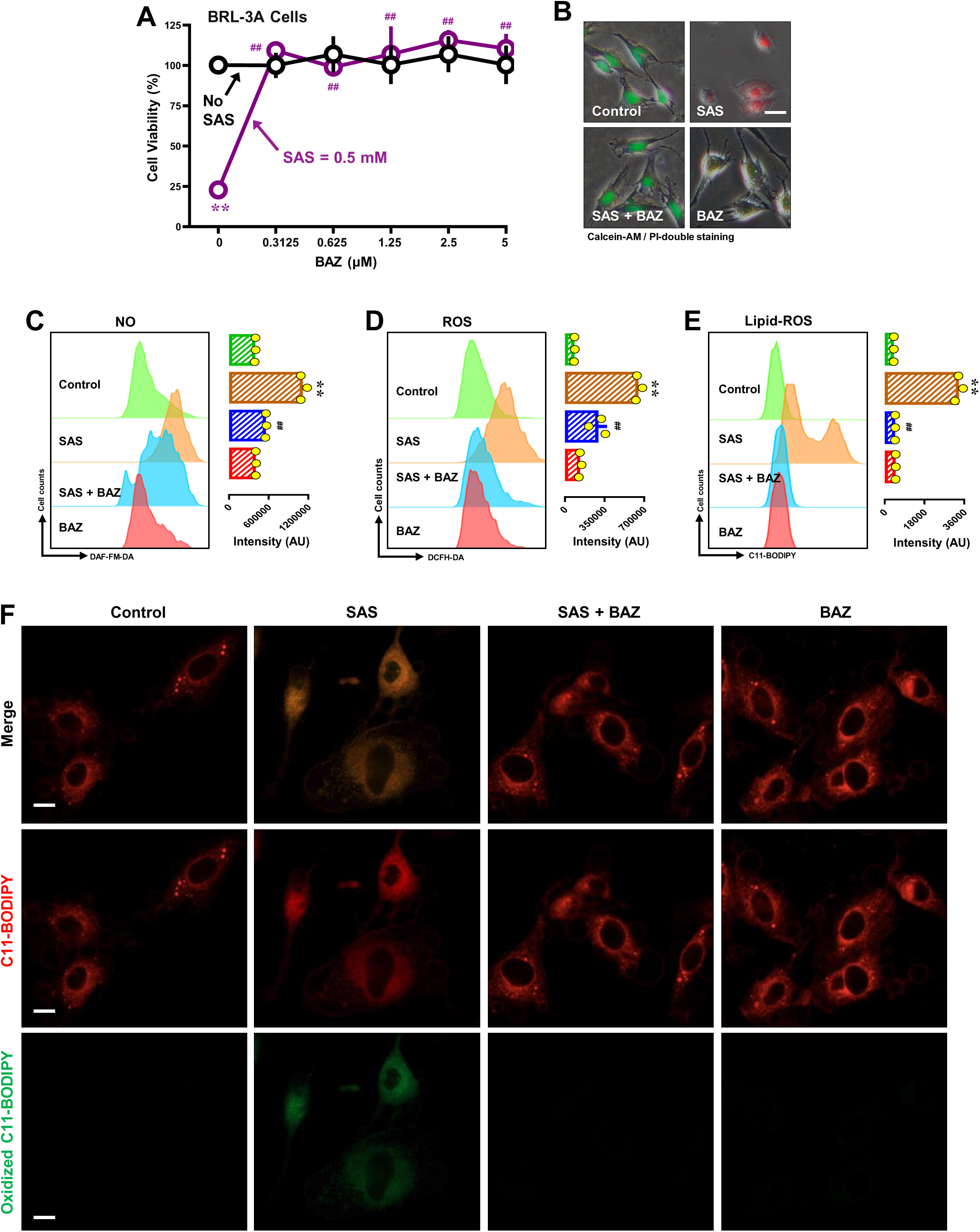
Effect of BAZ on SAS-induced ferroptosis and accumulation of NO, ROS and lipid-ROS in BRL-3A cells. **A, B**. Protective effect of BAZ against SAS-induced cytotoxicity. In **A**, cells were treated with SAS (0.5 mM) ± BAZ (0.15625, 0.3125, 0.625, 1.25, 2.5 and 5 µM) for 24 h, and cell viability was determined by MTT assay (n = 5). In **B**, cells were treated with SAS (0.5 mM) ± BAZ (2.5 µM) for 24 h, and then fluorescent images of Calcein-AM/PI-stained cells were captured (green for live cells, and red for dead cells; scale bar = 100 μm). **C, D, E.** Abrogation by BAZ of SAS-induced accumulation of cellular NO (**C**), ROS (**D**) and lipid-ROS (**E**). Cells were treated with SAS (0.5 mM) ± BAZ (2.5 µM) for 8 h, and then subjected to analytical flow cytometry. The left panels are the histograms, and the right panels are the quantitative values (n = 3). **F**. Abrogation by BAZ of SAS-induced accumulation of cellular lipid-ROS (confocal microscopy, scale bar = 100 μm). Quantitative data are presented as mean ± SD. ** or ^##^ *P* < 0.01; n.s., not significant.

**Supplementary Fig. S10.**
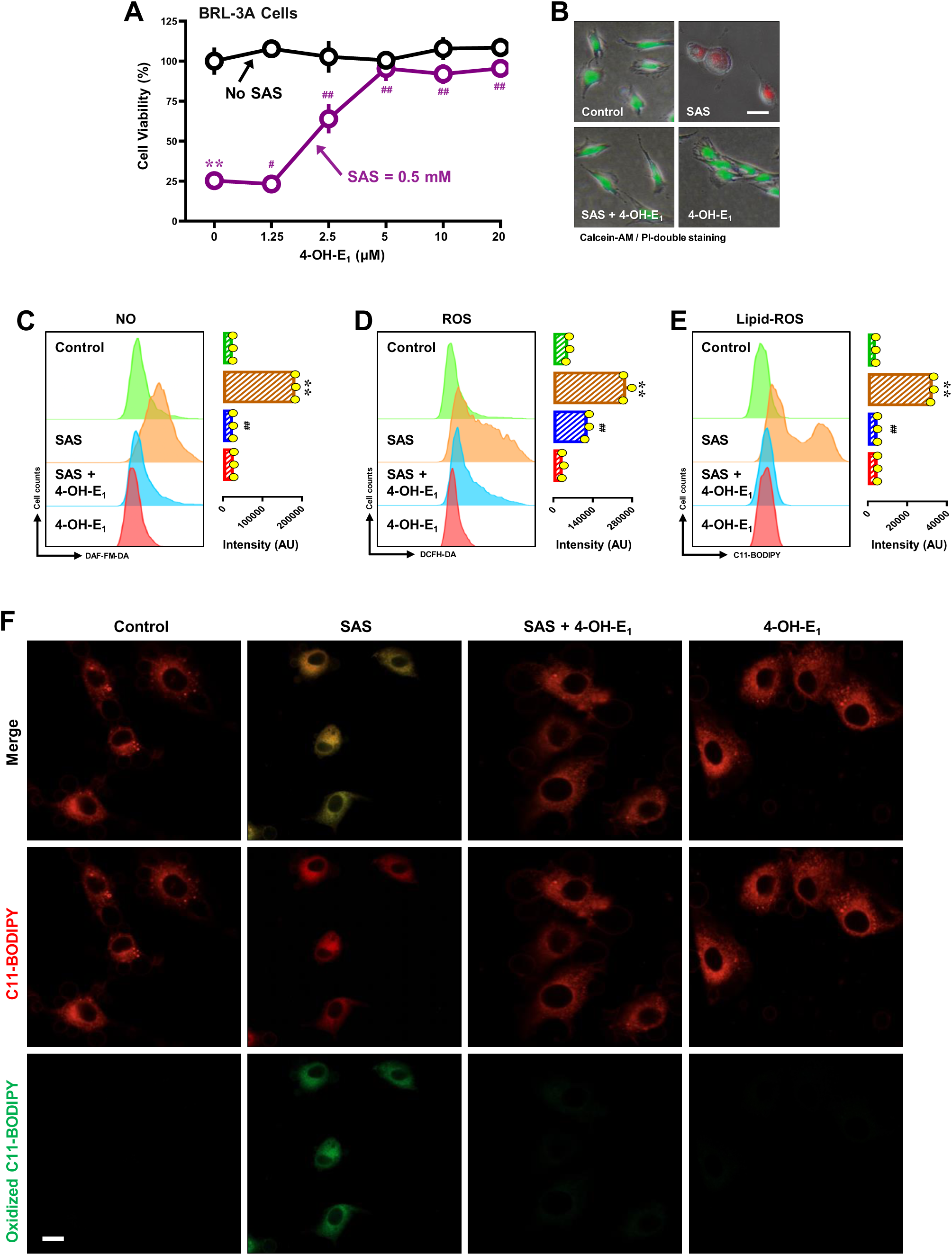
Effect of 4-OH-E_1_ on SAS-induced ferroptosis and accumulation of NO, ROS and lipid-ROS in BRL-3A cells. **A, B**. Protective effect of 4-OH-E_1_ against SAS-induced cytotoxicity. In **A**, cells were treated with SAS (0.5 mM) ± 4-OH-E_1_ (1.25, 2.5, 5, 10 and 20 µM) for 24 h, and cell viability was determined by MTT assay (n = 5). In **B**, cells were treated with SAS (0.5 mM) ± 4-OH-E_1_ (5 µM) for 24 h, and then fluorescent images of Calcein-AM/PI-stained cells were captured (green for live cells, and red for dead cells; scale bar = 100 μm). **C, D, E.** Abrogation by 4-OH-E_1_ of SAS-induced accumulation of cellular NO (**C**), ROS (**D**) and lipid-ROS (**E**). Cells were treated with SAS (0.5 mM) ± 4-OH-E_1_ (5 µM) for 8 h, and then subjected to analytical flow cytometry. The left panels are the histograms, and the right panels are the quantitative values (n = 3). **F**. Abrogation by 4-OH-E_1_ of SAS-induced accumulation of cellular lipid-ROS (confocal microscopy, scale bar = 100 μm) in BRL-3A cells. Quantitative data are presented as mean ± SD. ** or ^##^ *P* < 0.01; n.s., not significant.

**Supplementary Fig. S11.**
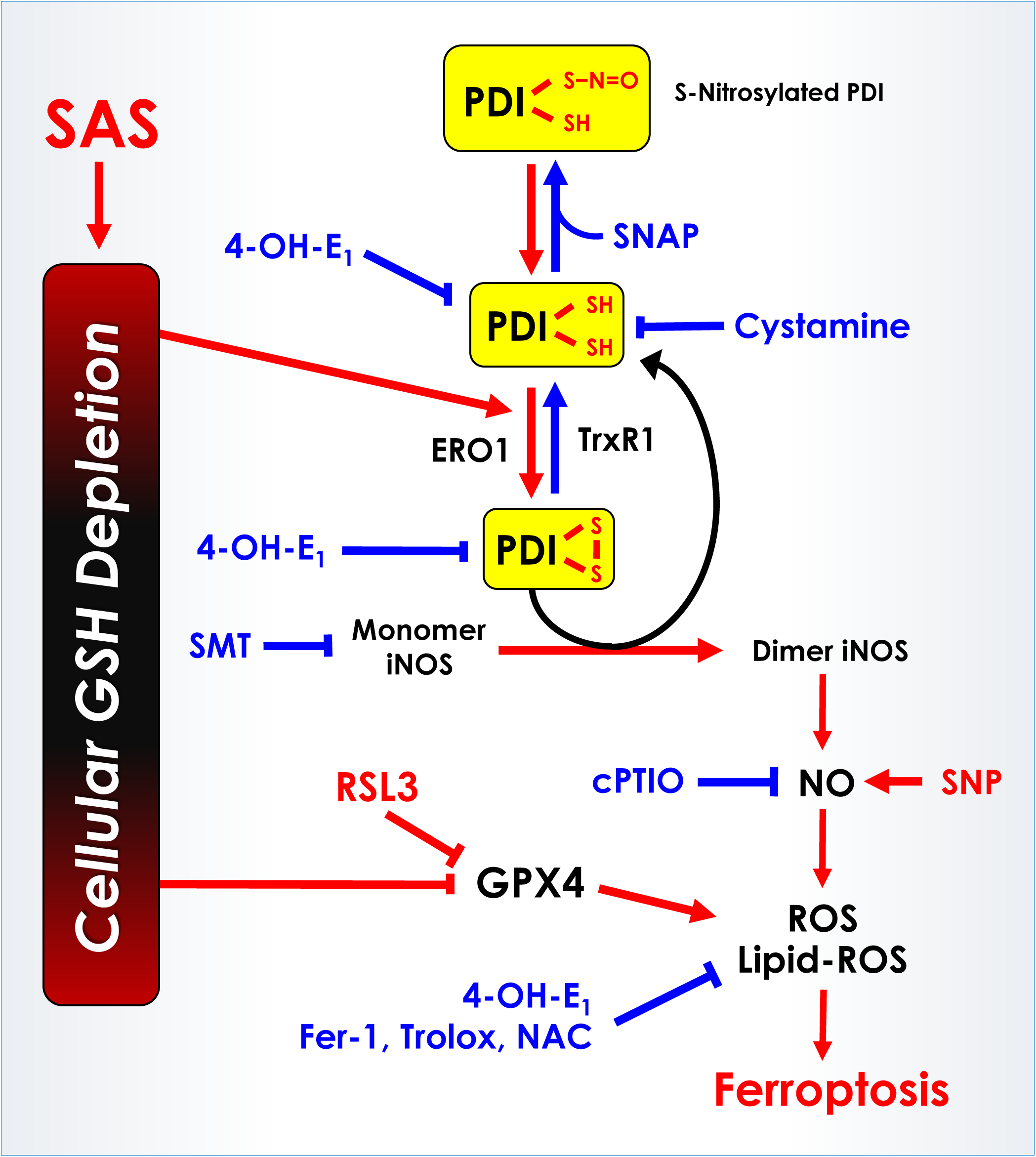
Schematic depiction of the role of PDI–iNOS/NO pathway in mediating SAS-induced ferroptosis. For details, please refer to the **DISCUSSION** section.

## Notes

**Conflict of interest:** All authors declare no conflict of interest.

**Funding:** This study was supported by research grants from Shenzhen Peacock Plan (No. KQTD2016053117035204), Shenzhen Key Laboratory of Steroid Drug Discovery and Development (No. ZDSYS20190902093417963), the National Natural Science Foundation of China (No. 81630096), Shenzhen Bay Laboratory (No. SZBL2019062801007).

### Competing Interest Statement

The authors have declared no competing interest.

### Summary of Updates

Nothing of the manuscript was changed from the earlier version, except that there was an error in the order of the authors in the online ABSTRACT page which has now been corrected so that all aurthors are listed in the same order as they appear in the manuscript. We apologize for the minor error.

